# Intra-membrane client recognition potentiates the chaperone functions of Calnexin

**DOI:** 10.1101/2022.03.07.483232

**Authors:** Nicolas Bloemeke, Kevin Meighen-Berger, Manuel Hitzenberger, Nina C. Bach, Marina Parr, Joao P.L. Coelho, Dmitrij Frishman, Martin Zacharias, Stephan A. Sieber, Matthias J. Feige

**Author notes:** Corresponding author: Matthias J. Feige, Technical University of Munich, Department of Chemistry and Institute for Advanced Study, Lichtenbergstr. 4, 85748 Garching, Germany.

## Abstract

One third of the human proteome are membrane proteins. They are particularly vulnerable to misfolding, often requiring assistance by molecular chaperones. Calnexin (CNX), one of the most abundant ER chaperones, plays an important role in membrane protein biogenesis and engages clients via its sugar-binding lectin domain.

Using mass spectrometric analyses, we show that Calnexin (CNX) interacts with a large number of non-glycosylated membrane proteins, suggesting additional binding modes. We find that misfolded membrane proteins are preferentially bound by CNX and that CNX uses its single transmembrane domain (TMD) for client recognition. Combining experimental and computational approaches, we systematically dissect signatures for intramembrane client recognition by CNX and identify sequence motifs within the CNX TMD region that mediate client binding. Building on this, we show that intramembrane client binding potentiates the chaperone functions of CNX.

Together, this study reveals a widespread role of CNX client recognition in the lipid bilayer, which synergizes with its established lectin-based substrate binding. Molecular chaperones thus can combine different interaction modes to support the biogenesis of the diverse eukaryotic membrane proteome.

## Introduction

Integral membrane proteins (IMPs) play essential roles in biology, including transporting molecules and signals across lipid bilayers, functioning as metabolic enzymes, and mediating cell-cell interactions. Around one third of all human genes code for membrane proteins (Fagerberg *et al*, 2010). In eukaryotic cells, the biosynthesis of both IMPs and soluble secretory pathway proteins occurs at the endoplasmic reticulum (ER) (Shao & Hegde, 2011). IMPs, however, often pose more complex challenges to the ER folding and quality control system than soluble proteins since the formation of their native state requires multiple topologically distinct folding events (Marinko *et al*, 2019). These include correct structuring of soluble domains on both the ER lumenal and cytosolic side and the proper lipid bilayer integration often including intra- and intermolecular assembly of transmembrane domain (TMDs) (Hegde & Keenan, 2021; O’Keefe *et al*, 2021). Further complicating these processes is the fact that TMDs of multipass membrane proteins are highly diverse in nature. They often contain polar residues, breaks or kinks and may exhibit flexibility in the membrane which gives rise to complex membrane integration and folding pathways (Feige & Hendershot, 2013; Hessa *et al*, 2005; Kauko *et al*, 2010; Lu *et al*, 2000; Ota *et al*, 1998; Sadlish *et al*, 2005). Notwithstanding, these deviations from ideal hydrophobic membrane anchors allow membrane proteins to fulfill their wide functional repertoire including transport of hydrophilic molecules through the lipid bilayer and specific recognition events in an apolar environment (Illergard *et al*, 2011). At the same time, they render IMPs vulnerable to incorrect folding and assembly in the lipid bilayer. This is demonstrated by the many severe human pathologies that are caused by membrane protein misfolding (Guerriero & Brodsky, 2012; Marinko *et al*., 2019; Partridge *et al*, 2004). Intramembrane chaperones and quality control factors that can efficiently recognize and monitor the folding status of IMPs directly within the lipid bilayer to support folding and detect misfolding are thus a prerequisite for protein homeostasis of any eukayotic cell.

One of the first chaperones identified to be involved in membrane protein folding was Calnexin (CNX) (Anderson & Cresswell, 1994; Hammond & Helenius, 1994; Jackson *et al*, 1994). CNX, which is tethered to the ER membrane via its single TMD, plays a crucial role in glycoprotein folding and transiently interacts with a wide range of newly synthesized proteins that transit the ER. As a part of the ER quality control system, CNX prevents incompletely folded substrates from leaving the ER and recruits co-chaperones that accelerate slow folding reactions including disulfide bond formation (Hebert & Molinari, 2012). The marked preference of CNX for monoglucosylated oligosaccharides on its clients results from its ER-lumenal lectin domain (Hebert *et al*, 1995). A highly similar lectin domain is found within Calreticulin (CRT), the soluble homologue of CNX in the ER (Kozlov *et al*, 2010b). Despite their almost identical lectin domains, genetic deletions of CNX and CRT have different effects. Whereas knockout cell lines of either protein are viable, organism develelopment is compromised. CRT deletion in mice leads to failures in heart development and prenatal lethality (Mesaeli *et al*, 1999). In contrast, deletion of CNX strongly affects nerve fibers and causes early postnatal death (Denzel *et al*, 2002). A major discriminating feature between CRT and CNX is the TMD of CNX. Previous work has found that anchoring CRT in the membrane through fusion with the TMD of CNX can alter the spectrum of proteins associated with CRT to be similar to that of CNX (Danilczyk *et al*, 2000; Wada *et al*, 1995). Although a simple explanation for functional differences may thus be the different localization of CNX and CRT, in the membrane versus ER-lumenal, this cannot account for many observations concerning the client binding of CNX. CNX directly interacts with the TM region of MHCI (Margolese *et al*, 1993). Furthermore, CNX recognizes and binds to truncated IMP substrates that lack one or more TM segments – presumably because they are recognized as misfolded and/or unassembled. Interestingly, in several of these studies CNX:substrate binding occurred in a glycan-independent manner (Cannon & Cresswell, 2001; Coelho *et al*, 2019; Fontanini *et al*, 2005; Li *et al*, 2010; Swanton *et al*, 2003; Wanamaker & Green, 2005). These and numerous other studies indicate that the CNX TMD is more than just a simple membrane anchor and potentially adds an essential function for intramembrane client recognition. Despite the abundance and key role of CNX in ER protein folding, no detailed insights are available into a potential intramembrane client recognition by CNX or its biological impact.

Here, we show that the substrate spectrum of CNX contains a large number of non-glycosylated membrane proteins and provide an in-depth analysis of intramembrane client recognition by CNX. Our study defines features within the TMD of CNX, as well as within its clients, that allow CNX to bind its substrates in the lipid bilayer. We further identify a motif within the CNX TMD that is required for efficient client binding. This allows us to reveal a protective role of intramembrane client binding for CNX substrates. Together, these structural, mechanistic, and systematic analyses provide a comprehensive understanding of intramembrane substrate recognition by the molecular chaperone CNX. Molecular chaperones can thus combine different binding modes to safeguard the biosynthesis of membrane proteins in the ER.

## Results

### Calnexin interacts with non-glycosylated membrane proteins in a chaperone-like manner

No systematic analysis on glycan-independent recognition of membrane protein clients by CNX exists yet. We thus decided to perform a global analysis of the CNX interactome. Toward this end we established CNX knockout cells and complemented these with a CNX construct with a C-terminal ALFA-tag (Fig 1A). The ALFA-tag is lysine-free (Götzke *et al*, 2019) and thus allows for the uncompromised use of lysine-crosslinkers, like DSSO, in co-immunoprecipitation experiments to stabilize transient CNX:client complexes. CNX itself has a large number of well-dispersed lysines which should allow for *in situ* DSSO-crosslinking (Fig 1B). Indeed, DSSO crosslinking led to many CNX crosslinks that could be immunoprecipitated using ALFA-tag nanobodies (Appendix Fig S1A). Mass spectrometry on the DSSO-crosslinked interactome of CNX revealed known ER-lumenal interactors like EDEM1, ERp29 and PDIA3, supporting the validity of our approach (Fig 1C). It furthermore revealed that approximately half of the interacting proteins were membrane proteins. Of note, among those, 44% were predicted to be not glycosylated (Fig 1C, Appendix Fig S1B). These include a small number of known functional interactors of CNX (e.g. the oxidoreductase TMX1 and the translocon subunit Sec63, Appendix Fig S1B) but most of the interactors are likely to be CNX clients.

**Figure 1.**
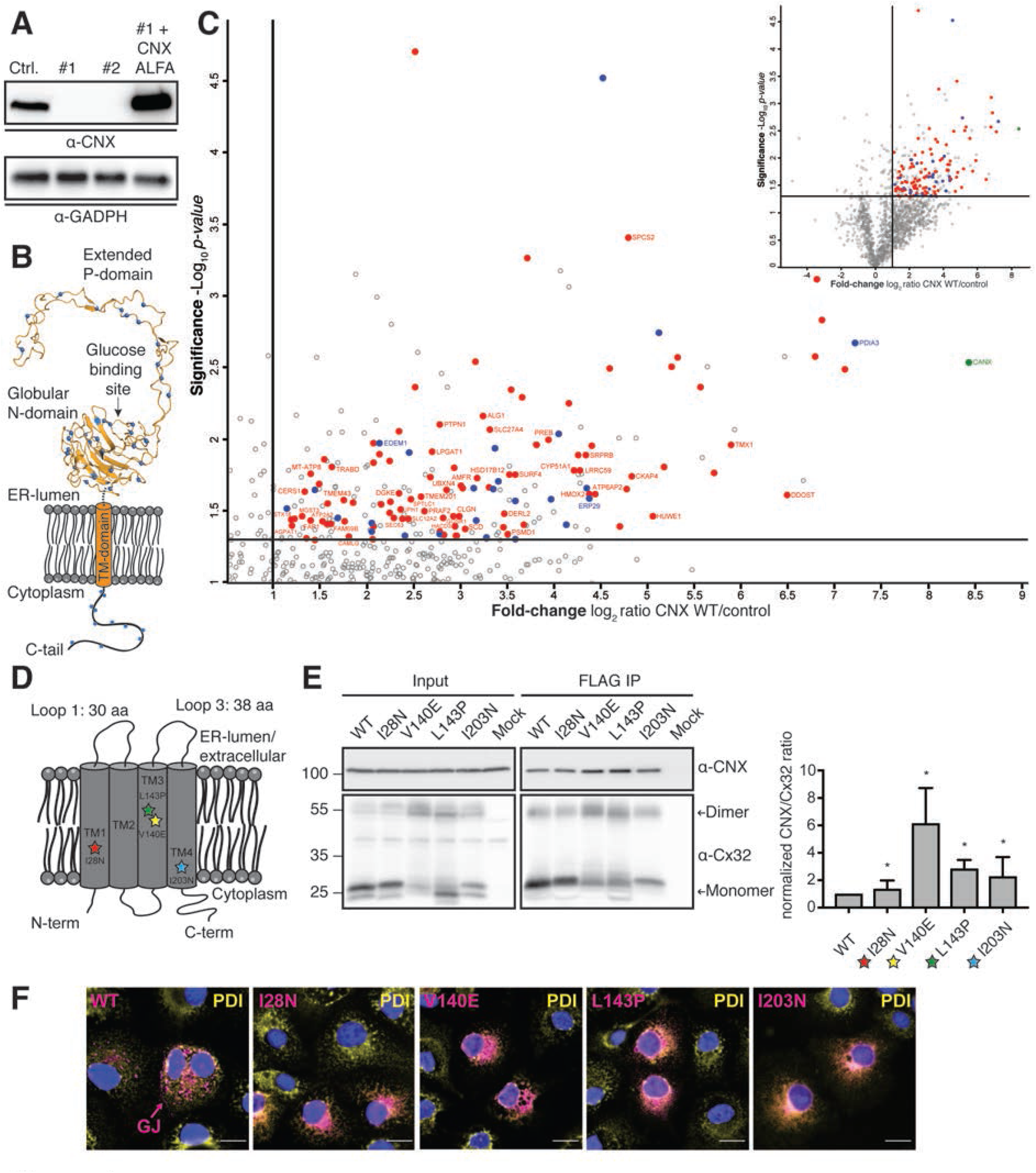
Calnexin interacts with non-glycosylated membrane proteins in a chaperone-like manner. A Immunoblot analysis of overexpressed and ALFA-tagged CNX in CNX knockout cell lines clones #1 and #2, which were used for mass spectrometry. Endogenous CNX level of wildtype HEK293T cells is shown as a control. B Schematic of CNX showing its lysine residues as blue dots. Lysines can be crosslinked by DSSO. The exact location of lysine residues is pictured in the crystal structure of the lumenal domain of CNX whereas the position of lysines in the cytoplasmic tail (C-tail) is an approximation. C Volcano plots derived from LC-MS/MS analysis of ALFA-tagged CNX, immunoprecipitated in 1% NP-40 buffer from transfected CNX deficient HEK293T cells after DSSO crosslinking and compared to empty vector (EV) control co-IPs. Depicted is an enlarged section of the associated complete volcano plot in the upper right. Among the significantly enriched proteins, annotations are Uniprot gene names. CNX is shown in green. ER proteins with the GO-term annotation “*endoplasmic reticulum”*, which are localized to the ER lumen, are highlighted in blue and known CNX interaction partners are indicated with their names. Membrane proteins with the GO-term annotation “*integral to membrane”* are highlighted in red. For non-glycosylated (according to Uniprot) integral membrane proteins, their name is additionally given. Integral membrane proteins not classified by the previously mentioned GO term annotation but possessing a TM domain were manually added to the interactome after additional analysis with Phobius (Käll *et al*., 2004). Cut-off values (solid lines) in the volcano plot were defined as log_2_ = >1 (2-fold enrichment) and -log_10_ = (P-value) > 1.3 (P<0.05). D Schematic of Cx32 structure and its orientation within the membrane. Positions of mutations analyzed in this study are indicated with stars. The overall topology and loop lengths were obtained from the UniProt server (human GJB1 gene). E Co-immunoprecipitations of endogenous CNX with FLAG-tagged Cx32 WT and its mutants including quantification and statistical analysis. Constructs were expressed in HEK293T cells. One representative immunoblot is shown. Monomers and dimers of Cx32 are indicated with arrows. All samples were normalized to WT (mean ± SEM, N ≥ 4, *P value < 0.05, two-tailed Student’s t tests). F Immunofluorescence images of Cx32 and its mutants. COS-7 cells were transiently transfected with the indicated FLAG-tagged Cx32 constructs and immunofluorescence microscopy was performed using anti PDI (yellow) as an ER marker. Detection of Cx32 was performed using anti FLAG antibodies and suitable labeled secondary antibodies (magenta). Nuclei were stained with DAPI (blue). All three channels are overlayed. Images are representative of cells from at least three different biological replicates. GJ denotes gap junctions observed in the cells transfected with WT Cx32. Scale bars correspond to 20 µm.

The large number of membrane proteins that interact with CNX, which are predicted to be non-glycosylated, suggests that this kind of interaction is very common. Furthermore, proteome-wide analyses in CNX k/o cells revealed reduced levels for several non-glycosylated membrane proteins (Appendix Fig S1C), implying a chaperone function of CNX for this protein class. Together, the interaction of CNX with non-glycosylated membrane proteins thus warrants further investigation.

To provide mechanistic insights into possible intra-membrane, glycan-independent client recognition and chaperoning processes by CNX, we used Connexin 32 (Cx32) as a first model protein, which a recent study from us has revealed to interact with CNX (Coelho *et al*., 2019). Cx32 is a tetra-spanning integral membrane protein (Fig 1D) that forms homo-hexameric connexons. Two of these connexons embedded in different membranes can dock onto each other to form a gap junction channel (Maeda *et al*, 2009; Pantano *et al*, 2008). Mutations in Cx32 cause X-linked Charcot-Marie-Tooth disease (CMTX), a common genetic disorders of the peripheral nervous system (Scherer & Kleopa, 2012). Cx32 is not glycosylated (Appendix Fig S1D) and possesses only short ER-lumenal loops (Fig 1D). The observed CNX binding can thus neither rely on recognition of sugar moieties nor on binding of large ER-lumenal domains by the CNX lectin domain. Taken together, we considered Cx32 a relevant starting point towards defining possible intra-membrane client binding by CNX.

To investigate the nature of the Cx32:CNX interaction in more detail, we expressed FLAG-tagged Cx32 in human HEK293T cells and analyzed its interactions with endogenous CNX in co-immunoprecipitation experiments. These confirmed interaction of Cx32 with CNX (Fig 1E) and revealed interaction of CNX with monomeric and dimeric Cx32 (Appendix Fig S1E). If CNX indeed acted as a chaperone on Cx32, one would expect a preferential interaction with misfolded variants of the client. We thus proceeded to study four disease-causing mutants of Cx32 (Bone *et al*, 1997; Kleopa *et al*, 2006; Rouger *et al*, 1997). All of these contain mutations in their TMDs (Fig 1D), but are properly integrated into the lipid bilayer (Appendix Fig S1D). In contrast to Cx32 wild type (Cx32-WT), all mutants were retained in the ER, arguing for misfolding and recognition by the ER quality control system (Fig 1F). Strikingly, all of the mutants showed a significant increase in interaction with CNX, which was up to ∼6-fold stronger than the interaction with Cx32-WT (Fig 1E). Taken together, our data show that CNX interacts with a large number of non-glycosylated membrane proteins, is important for their normal cellular levels and that interactions occur in a chaperone-like manner.

### The transmembrane domain of Calnexin binds clients in the membrane

Our data suggest that CNX can recognize misfolded Cx32 in the plane of the membrane where the mutations are located. To further test this hypothesis, we individually fused each of the four Cx32 TMDs to an antibody light chain constant domain (C_L_) (Fig 2A). A related system was established previously to assess chaperone:client interactions for soluble proteins in a systematic manner (Behnke *et al*, 2016; Feige & Hendershot, 2013). By using a glycosylation reporter site downstream of the TMD we could verify that the majority of each TMD segment was integrated into the ER membrane. The only exception was TMD2 (Appendix Fig S2A), which did not integrate properly, in agreement with a recent study (Coelho *et al*., 2019) and the predicted membrane integration potentials (Fig 2A). TMD2 was thus excluded from further analyses. As a small portion of the TMD3 and TMD4 constructs could also enter the ER, we replaced the sugar-accepting Asn residue downstream of each TMD by a Gln residue, to exclusively monitor glycan-independet interaction with CNX. Using this system, we probed if CNX could bind individual TMD segments and if so, which of the TMD segments from Cx32 was bound by CNX. Interaction was observed for all three TMD segments and among those, by far the strongest interaction was observed with TMD1 (Fig 2B). Introducing mutations into the individual TMD segments compromised their proper membrane integration (Appendix Fig S2B and C), precluding further direct analyses on the effects of mutations on binding. Collectively, our data show that CNX can differentially bind individual TMD segments of a non-glycosylated client.

**Fig 2.**
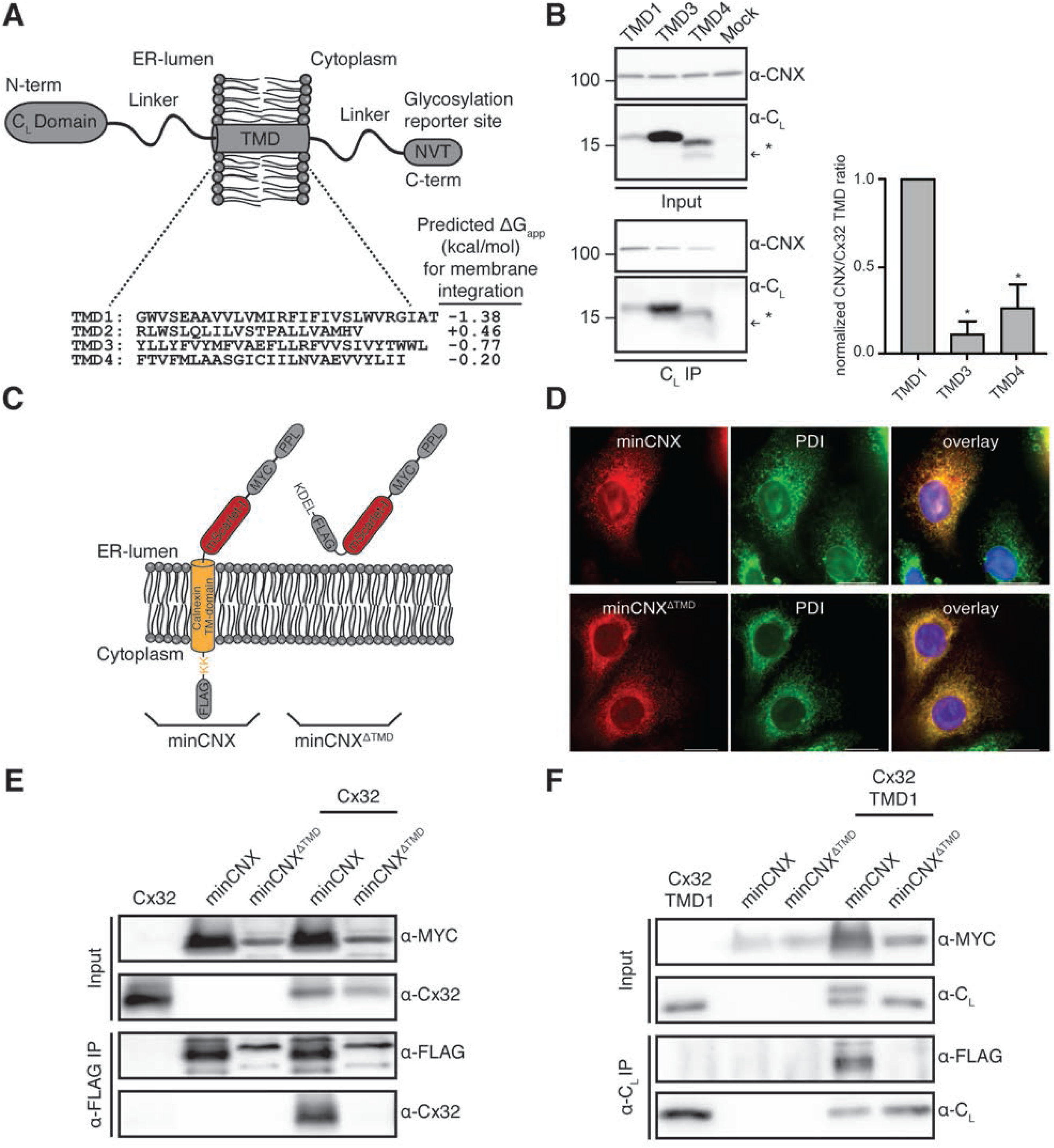
The transmembrane domain of Calnexin binds clients in the membrane. A Schematic of the reporter construct, where an antibody C_L_ domain was fused to seuences of interest, to assess CNX interactions with individual Cx32 TMD segments. The four WT sequences of the Cx32-TMDs are shown. Predicted membrane integration energies are given using DGPred (Hessa *et al*., 2007). Sequences of TMD1 and TMD3 were inverted in the respective constructs to reflect the correct orientation within Cx32. B Co-immunoprecipitations of endogenous CNX with the different C_L_-TMD constructs (NVT glycosylation site mutated to QVT) including quantification and statistical analysis. One representative immunoblot is shown. The asterisk (*) denotes a fraction of cleaved species as observed previously for similar constructs (Behnke *et al*., 2016). All samples were normalized to the construct containing TMD1 (mean ± SEM, N ≥ 3, *P value < 0.05, two-tailed Student’s t tests). C Schematic of the minimal CNX construct (minCNX). This consists of a preprolactin (PPL) ER import sequence, followed by an N-terminal MYC tag, the monomeric red fluorescent protein mScarlet-I, the TMD segment of CNX including an endogenous double lysine motif downstream of the TMD region and a C-terminal FLAG-tag. Flexible linker regions connect the individual components. Parts derived from CNX are shown in orange. A control construct, minCNX^ΔTMD^, is lacking the entire CNX TMD segment. A C-terminal KDEL sequence was included for ER retention. D COS-7 cells were transiently transfected with the indicated constructs and immunofluorescence microscopy was performed using anti PDI (green) as an ER marker. Detection of minCNX constructs was carried out by mScarlet-I fluorescence (red). Nuclei were stained with DAPI (blue). Images are representative of cells from at least three different biological replicates. Scale bars correspond to 20 µm. E Interaction of wild type Cx32 with the minCNX system. MinCNX constructs (see (C)) were co-transfected with Cx32 WT into HEK293T cells. Interaction between the proteins was analyzed by co-immunoprecipitation experiments followed by immunoblotting. F Interaction of Cx32-TMD1 with the CNX TMD. Representative blots from co-immunoprecipitation and immunoblotting experiments from HEK293T cells co-transfected with the indicated Cx32-TMD1 and minCNX constructs including are shown.

Based on these findings, we next aimed at defining which structural elements of CNX were necessary and sufficient for this interaction. Toward this end, we designed a minimal CNX (minCNX) construct that only contained the CNX TMD and a few additional C-terminal residues. The ER-lumenal domain of CNX was replaced by the fluorescent protein mScarlet-I (Fig 2C). To maintain ER retention of the construct, an endogenous cytosolic di-lysine motif was left in place (Fig 2C, Appendix Fig S2C) (Jackson *et al*, 1990). ER-localization was confirmed by fluorescence microscopy (Fig 2d). Using this construct, we found that the CNX TMD was necessary and sufficient for binding to full-length Cx32 (Fig 2E). Further extending this finding, we could show that the first TMD of Cx32 was sufficient to bind to the minCNX construct (Fig 2F). In either case, an ER-retained control construct lacking the CNX TMD region (minCNX^ΔTMD^, Fig 2D, Appendix Fig S2E) did not bind to the different clients (Fig 2E and F). Thus, binding to the non-glycosylated TM clients occurs via the CNX TMD, independently of the CNX lectin domain or its C-terminal tail. Accordingly, the CNX TMD contains the relevant features for client recognition in the membrane, and this can be recapitulated with a single TMD derived from a client protein.

### Development of a tool to systematically assess intramembrane recognition processes

Our data show that CNX directly binds to individual TMD segments of its clients and that the CNX TMD is sufficient for binding to occur. This established a minimal system for client recognition by a chaperone in the membrane. Based on these findings, we next aimed to define which intramembrane features of its clients CNX recognizes. This would be a major step forward in our understanding of intra-membrane chaperones but is a difficult task to accomplish using (parts of) natural proteins, as these generally will possess complex sequences and structures which precludes unbiased analyses. We thus designed a client protein that allows for the systematic analysis of intramembrane recognition processes. Toward this end, we performed a multiple sequence alignment of 200 randomly selected human single pass plasma membrane proteins from the Membranome database (Lomize *et al*, 2018; Lomize *et al*, 2017). This gave rise to an average TMD of a transport-competent protein, which can thus be expected to lack major chaperone recognition sites. We termed this protein *minimal <u>con</u>sensus <u>mem</u>brane protein* (ConMem). The design of ConMem included a superfolder GFP (sfGFP) moiety for microscopic localization studies, a C-terminal HA-epitope tag, and two glycosylation sites to analyze the membrane topology and intracellular transport of the reporter (Fig 3A). In agreement with our assumption that, on average, TMD segments from single-pass cell surface TMD proteins are close to optimal, the most frequent amino acid at most positions turned out to be Leu (Fig 3A). TM Leu-zippers, however, have a strong self-assembly propensity that could compromise our analyses (Gurezka *et al*, 1999). We thus proceeded with the second most frequently occuring amino acids, which were also entirely free of unfavorable residues for a TM sequence (Hessa *et al*, 2007). The multiple sequence alignment resulted in a TMD of 26 amino acid in length in ConMem, which is in very good agreement with recent studies on average TMD lengths in the plasma membrane (Sharpe *et al*, 2010; Singh & Mittal, 2016). Interestingly, it was flanked by an N-terminal Pro-residue, breaking the helical TMD structure (Cordes *et al*, 2002), and a C-terminal Lys residue (Fig 3A, Appendix Fig S3). A C-terminal Lys will induce a type I orientation (von Heijne & Gavel, 1988), placing the Lys residue in the cytoplasm, which reflects the nature of our sequence set (76% of the proteins were type I). Individually mutating the first or second consensus glycosylation site in ConMem showed that it was exclusively modified at the first site, which confirms the predicted topology (Fig 3B). Enzymatic deglycosylation with EndoH (which only removes N-linked sugars not further modified further in the Golgi) or with PNGaseF (which removes all N-linked sugars) revealed that ConMem glycosylation was EndoH-resistant, arguing that it was able to traverse the Golgi as expected (Fig 3B). Localization to the plasma membrane was confirmed by microscopic studies (Fig 3C). Taken together, ConMem provided us with an ideal tool to systematically dissect intramembrane substrate recognition by CNX.

**Fig 3.**
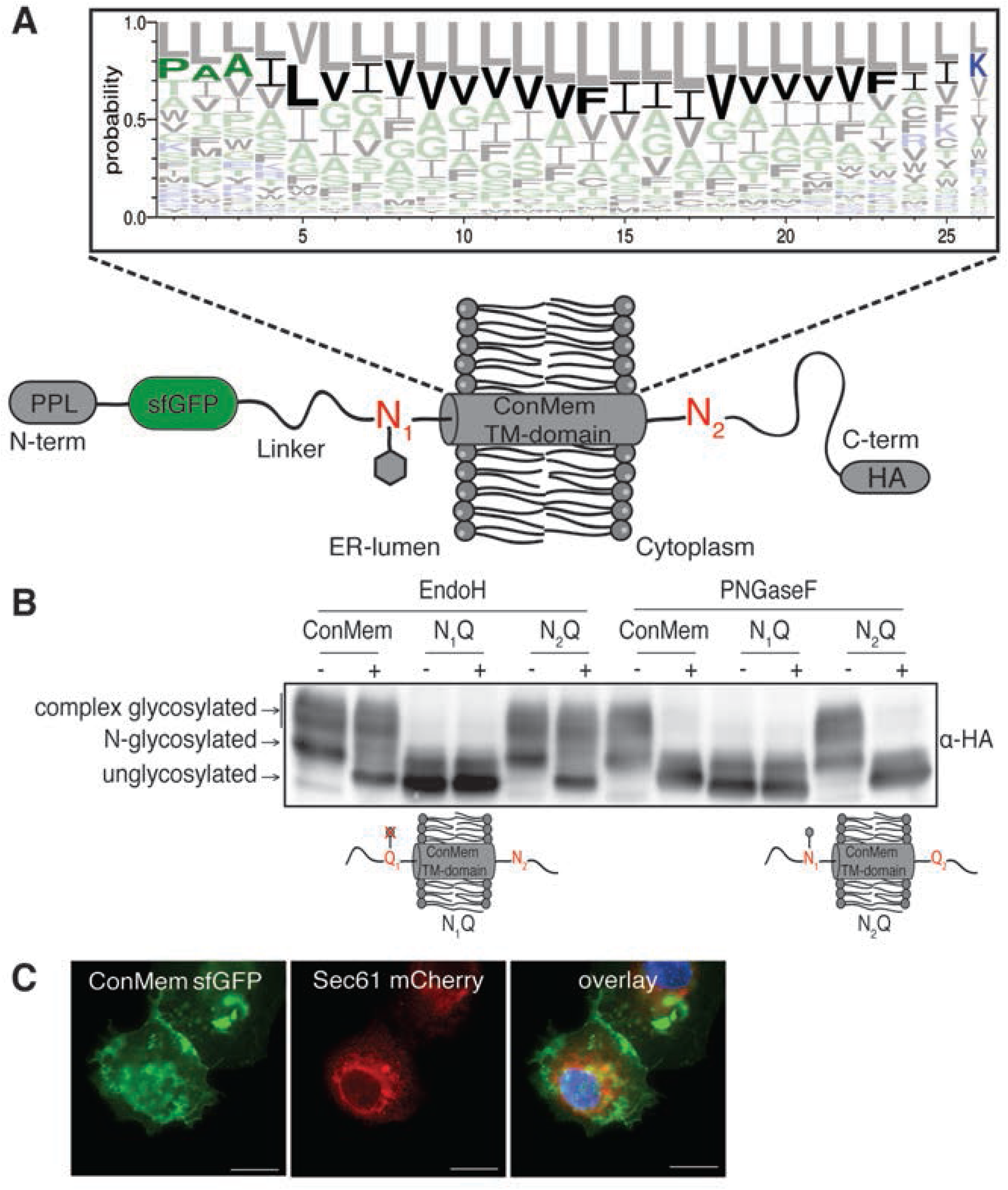
Design and validation of ConMem, a tool to query TM-domain recognition. A Schematic of the the minimal consensus membrane protein *ConMem* which contains a preprolactin (PPL) ER import sequence, a superfolder GFP (sfGFP), a TMD flanked by two individual NVT glycosylation motifs (N_1_ and N_2_) and a C-terminal HA-tag. Individual construct components are connected by flexible linker regions. As illustrated, in the predicted topology of ConMem, only the first NVT glycosylation site (N_1_) is accessible to the ER glycosylation machinery (grey hexagon). The TMD of ConMem was designed on the basis of a multiple sequence alignment of 200 human single pass TMD sequences as illustrated in the sequence logo. Hydrophilic amino acid residues are depicted in blue, neutral ones in green and hydrophobic residues in black. The predicted amino acid sequence with the second highest score was selected as the TMD consensus sequence for ConMem (bold). B HEK293T cells were transiently transfected with plasmids expressing ConMem or the indicated variants, where either the first (N_1_) or the second (N_2_) NVT glycosylation motif was ablated (N to Q mutation). Lysates were treated with or without EndoH or PNGaseF as indicated, and analyzed by immunoblotting. N-glycosylation occurs in the ER, complex glycosylation in the Golgi. C COS-7 cells were transfected with the indicated constructs and fluorescence microscopy was performed using Sec61 mCherry (red) as an ER marker. ConMem, detected by sfGFP fluorescence (green), localizes to the plasma membrane. Nuclei were stained with DAPI (blue). Images are representative of cells from at least three different biological replicates. Scale bars correspond to 20 µm.

### Defining intramembrane recognition motifs for Calnexin

To dissect features recognized by CNX in the membrane, we first replaced the central Val residue at position 13 in ConMem by all other 19 amino acids. Of note, for all of these substitutions, ConMem was still predicted to be stably integrated into the membrane (Fig 4A), which was experimentally confirmed for several constructs and different locations of polar amino acids (Appendix Fig S4A). Using this ConMem panel, we first established suitable conditions for co-immunoprecipitation experiments (Appendix Fig S4B) and then investigated interactions with CNX. Although ConMem containing an ER-lumenal glycosylation site bound stronger to CNX, significant binding was also observed without this site. This shows that glycosylation of ConMem increases binding to CNX but is not required for it to occur (Fig 4B), in agreement with our mass spectrometric analyses and the binding to Cx32 (Fig 1). No binding of sfGFP to CNX was observed (Appendix Fig S4C), neither did Cx32 bind to ConMem (Appendix Fig S4D), showing the specificity of the ConMem:CNX interactions. Together, this further corroborated binding of CNX to TM regions in the membrane. For all subsequent experiments, ConMem lacking its ER-lumenal glycosylation site was used to specifically investigate glycan-independent binding to CNX. Since CNX also bound to ConMem with a mostly an ideal TMD segment, this allowed to assess features increasing and decreasing binding to CNX. Using this approach revealed a highly distinct binding pattern for CNX to the 20 ConMem variants with the membrane-central amino acid exchanged (Fig 4A). For some, e.g. Arg, binding was significantly increased, whereas for others, e.g. Pro, binding was decreased (Fig 4C, Appendix Fig S4E). Arg introduces a polar residue into the membrane, whereas Pro acts as a TMD helix breaker. Based on these findings, we selected one amino acid where increased binding to CNX was observed if it was present in a central location in the ConMem TMD segment (Arg), and one that decreased binding (Pro) (Fig 4C). For these, we moved the mutation site through the entire ConMem TMD to analyze a possible positional dependency on CNX binding (Fig 4D). In each case, the central position 13 (out of 26 amino acids) showed the strongest effects, but replacements at other positions also influenced binding, with slightly different positional dependencies for the two selected amino acids (Fig 4D). In summary, these data show that CNX can differently recognize clients with changes of single amino acids in the membrane.

**Fig 4.**
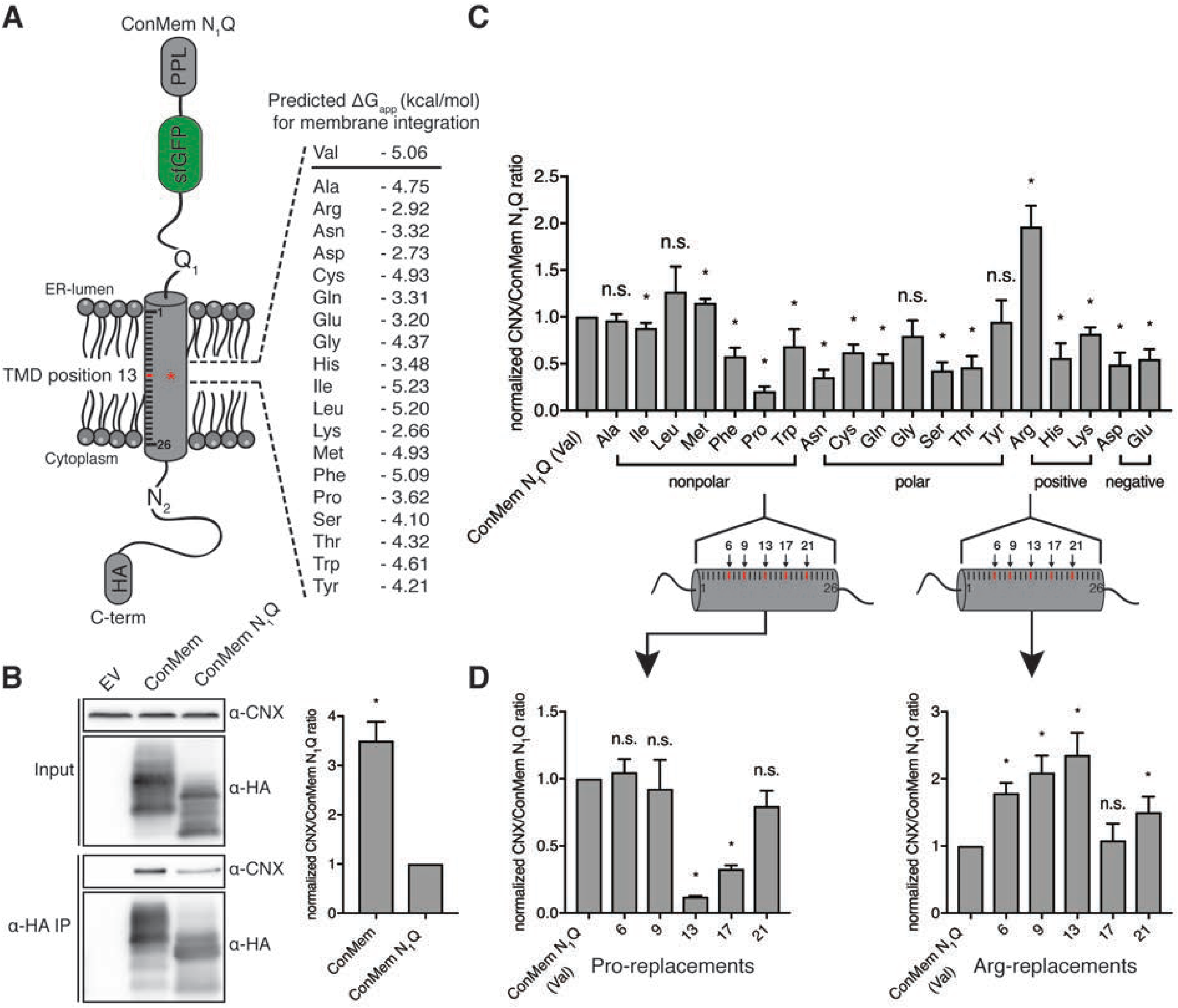
Defining intramembrane recognition motifs for Calnexin. A Schematic of ConMem N1Q where a central Val residue at position 13 within its TMD was replaced by all other 19 amino acids. Free energies for TMD insertion were predicted according to (Hessa *et al*., 2007). B Representative blots from co-immunoprecipitation experiments from HEK293T cells transfected with the indicated ConMem constructs with or without ER-lumenal glycosylation site (N1) (mean ± SEM, N ≥ 3, *P value < 0.05, two-tailed Student’s t tests). C Interaction of ConMem N1Q variants as described in (A) with endogenous CNX (mean ± SEM, N ≥ 5, *P value < 0.05, two-tailed Student’s t tests). D Same as in (C) only that selected amino acids (proline and arginine) where shifted through the ConMem N1Q TMD segment to five different positions as shown in the illustration, revealing positional binding dependencies (mean ± SEM, N ≥ 4, *P value < 0.05, two-tailed Student’s t tests). Individual values in (B-D) were normalized to ConMem N1Q (Val) values that were set to 1.

### A structural understanding of intramembrane Calnexin:client recognition

Having defined client-intrinsic binding patterns for CNX in the membrane, we next proceeded to analyze the features within the CNX TM segment that allow client binding to occur. Towards this end, we performed molecular dynamics simulations on either the CNX TM region together with the ConMem TMD or with the first TMD of Cx32 (Cx32-TMD1), which our data have shown to be interacting systems in cells (Fig 2 and 4). In agreement with these experimental findings, molecular dynamics simulations also showed interactions for both systems (Fig 5A). Of note, for ConMem, as well as for Cx32-TMD1, similar regions in the CNX TMD region were involved in the binding process. These regions involved a Tyr, a Thr and a Leu on the same face of the CNX TMD region (Fig 5A-C). To assess our computational predictions experimentally, we mutated these predicted interaction sites to presumably inert Val residues in a CNX construct that we expressed in mammalian cells. This construct was furnished with a V5 epitope tag for specific immunoprecipitation and overexpression was only slightly higher than the endogenous CNX level (Appendix Fig S5A). Strikingly, whereas ConMem and Cx32- TMD1 co-immunoprecipitated with V5-tagged CNX as expected, mutation of the YTL-motif, and even of only the Y and T residues to Val, significantly reduced interactions between CNX and ConMem (Fig 5D). This was even more pronounced for Cx32-TMD1 (Fig 5E), and when the YTL-motif was exchanged against Ala instead of Val, a similar reduction in binding was observed (Appendix Fig S5B). Furthermore, when we assessed two of the non-glycosylated TM protein interactors of CNX our mass spectrometry analysis had revealed (Fig 1C, Appendix Fig S1B) we also observed significant reduction in binding to CNX when the YTL-motif was mutated (Appendix Fig S5C). Together this argues that mutating the interaction site *per se*, but not the choice of the residues mutated to, accounts for the effects. Furthermore, these findings confirmed our simulation data and further pointed towards a conserved interaction motif in CNX, necessary for client binding in the membrane. Of note, when ConMem with its ER-lumenal glycosylation site was used, mutations in the CNX TMD region did not significantly affect binding, arguin that for this glycosylated client with a well-behaving TMD, binding via the lectin domain is dominant (Fig 5F). The importance of the intramembrane YTL-motif obtained with these model clients was corroborated when we investigated CNX interactions with full-length Cx32 (Fig 5G) as well as the Cx32 V140E mutant (Appendix Fig S5D).

**Fig 5.**
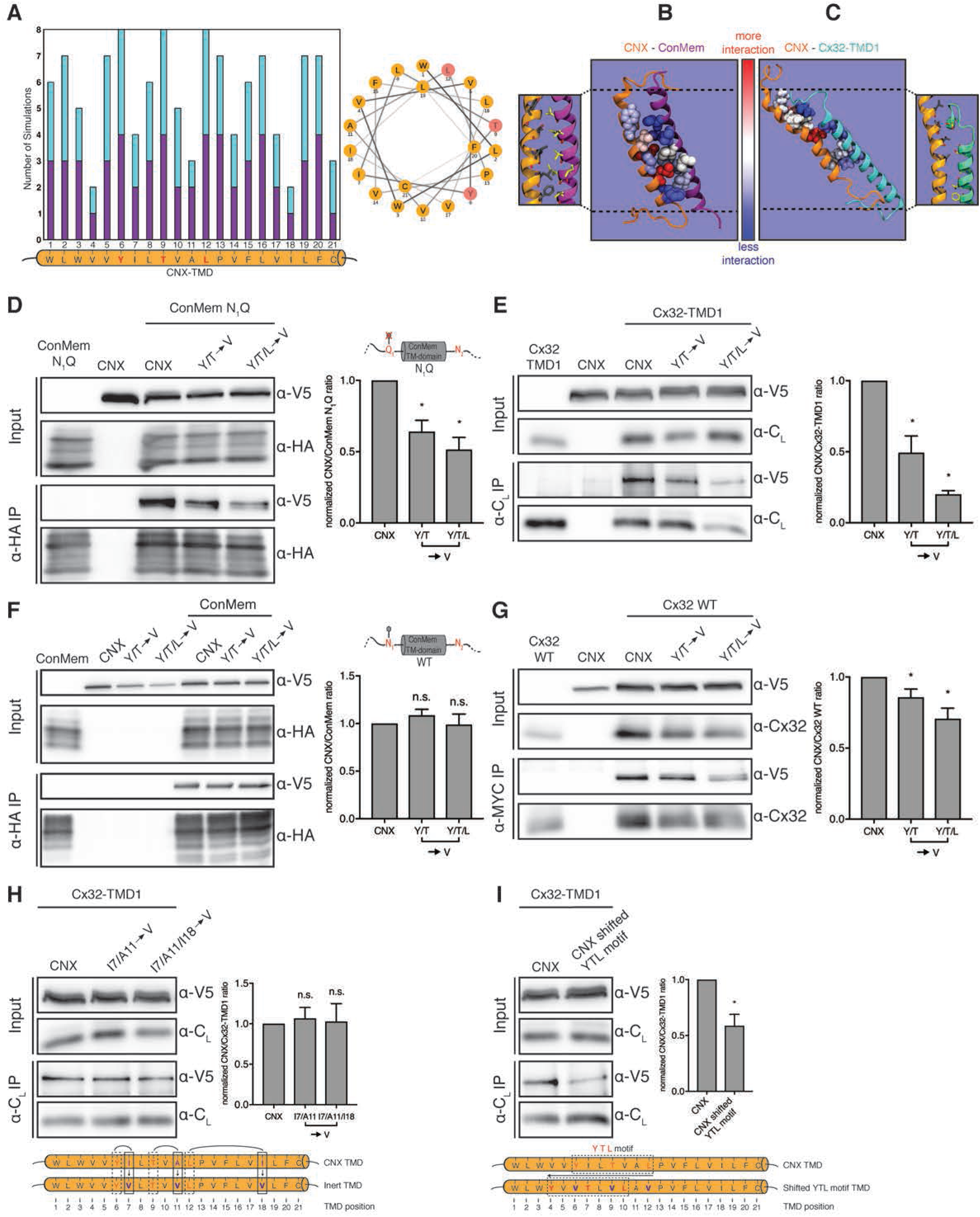
A structural analysis of intramembrane Calnexin:client recognition. A Meta analysis of MD simulations (N=8) where complex formation between CNX and Cx32-TMD1 or CNX and ConMem was observed. On the x-axis the CNX TMD residues obtained from UniProt are shown. The height of the bars corresponds to the number of simulations in which a specific CNX TMD residue interacted with the substrate helix CX32-TMD1 (turquoise) or ConMem (violet). Only CNX TMD residues Y6, T9 and L12 interacted with CX32-TMD1 or ConMem in all sampled complexes. B Depiction of residues exhibiting most interactions with the partner helix in simulations leading to complexes between CNX and ConMem. C Same as (B) but for the CNX-Cx32-TMD1 simulations. CNX is depicted in orange, while ConMem is shown in purple and Cx32-TMD1 in turquoise. CNX, Cx32-TMD1 or ConMem residues exhibiting most frequent interactions with residues on the opposite helix are colored red, while residues in dark blue formed less interactions. For greater clarity, only the top most interacting residues are shown here. D Interaction of non glycosylated ConMem N_1_Q constructs with V5-tagged CNX. In CNX variants, amino acids important for interaction as revealed in (A) were replaced by valine residues. HEK293T cells were transiently transfected with the indicated constructs and cell lysates and HA-immunoprecipitates were analyzed for HA-tagged ConMem N_1_Q and co-immunoprecipitating CNX mutants. One representative blot and quantifications are shown (mean ± SEM, N ≥ 9, *P value < 0.05, two-tailed Student’s t tests). E The same as in (D) for Cx32-TMD1 (mean ± SEM, N ≥ 6, *P value < 0.05, two-tailed Student’s t tests). F Analysis of mutations in the CNX TMD region and their effect on binding glycosylated ConMem. One representative blot and quantifications are shown (mean ± SEM, N ≥ 3, n.s.: not significant, two-tailed Student’s t tests). G The same as in (B) for Cx32 full-length (mean ± SEM, N ≥ 4, *P value < 0.05, two-tailed Student’s t tests). Interaction of Cx32-TMD1 with CNX mutants, where amino acids not predicted to be important for interaction as shown in (A) were exchanged by valine residues. HEK293T cells were transiently transfected with the indicated constructs and a representative blot is shown. One representative blot and quantifications are shown (mean ± SEM, N ≥ 9, n.s.: not significant, two-tailed Student’s t tests). I Shifting of the residues Y, T and L, most important vor CNX substrate interaction, and replacing these against valine residues, and the impact on interactions with Cx32-TMD1. HEK293T cells were transiently transfected with the indicated constructs. One representative blot and quantifications are shown (mean ± SEM, N ≥ 9, *P value < 0.05, two-tailed Student’s t tests).

To further verify the specificity of the binding motif we have identified within the CNX TMD, we selected three non-Val residues that had the lowest overall predicted client interactions from our simulations (an Ile (TMD position 7), an Ala (TMD position 11) and another Ile (TMD position 18), Fig 5A). Replacing either the first two or all three of these three residues by Val did not affect binding of CNX to its client Cx32-TMD1 (Fig 5H), further supporting our molecular interpretation of CNX client binding in the membrane. In contrast, when we shifted the identified YTL-motif N-terminally within the CNX TMD, this again significantly reduced binding to Cx32-TMD1 (Fig 5I), revealing positional specificity in the recognition process. Our combined computational and experimental approach thus revealed a molecular recognition motif within the CNX TMD that CNX uses to bind clients in the lipid bilayer.

### Biological functions of intramembrane client recognition by Calnexin

Our comprehensive analyses provided us with detailed molecular insights into CNX:client recognition in the membrane. Using these insights, we were able to generate a CNX variant that was compromised in client binding within the membrane while still having a functional lectin domain (Fig 5). To define functional consequences of weakening intra-membrane client binding we used rhodopsin (Rho) as a model protein, as its biogenesis is highly dependent on CNX (Rosenbaum *et al*, 2006). To avoid confounding effects by endogenous CNX, we used our CNX knockout cell line (Fig 1A). CNX knockout did not cause detectable ER stress and neither did the re-expression of CNX or its mutant compromised in intramembrane client binding (Appendix Fig S6A). Furthermore, both CNX variants showed normal ER localization in the knockout cells (Appendix Fig S6B), which together are prerequisites for unbiased analyses. We first analyzed interaction of Rho with either reconstituted wt CNX or the mutant lacking the intramembrane YTL-motif. In agreement with our data on a panel of other model proteins (Fig 5, Appendix Fig S5), Rho binding to the CNX mutant was significantly reduced (Fig 6A), qualifying Rho as another candidate of the intramembrane client binding by CNX. In agreement with this finding, Rho as a strong CNX client was hardly detectable if CNX was completely absent (Fig 6A). Building on these insights, we assessed the effects of CNX on Rho stability in cells. Strikingly, Rho degradation was significantly accelerated in CNX knockout cells complemented with the YTL-motif mutant in comparison to wt CNX (Fig 6B), revealing a protective role of intramembrane substrate binding by CNX for a labile client. This protective role of intramembrane client binding appeared to work syngeristically with lectin-based client binding as the complete absence of CNX had an even stronger effect on Rho degradation (Fig 6B). Thus, CNX intramembrane client binding synergizes with lectin-based binding to chaperone labile clients.

**Fig 6.**
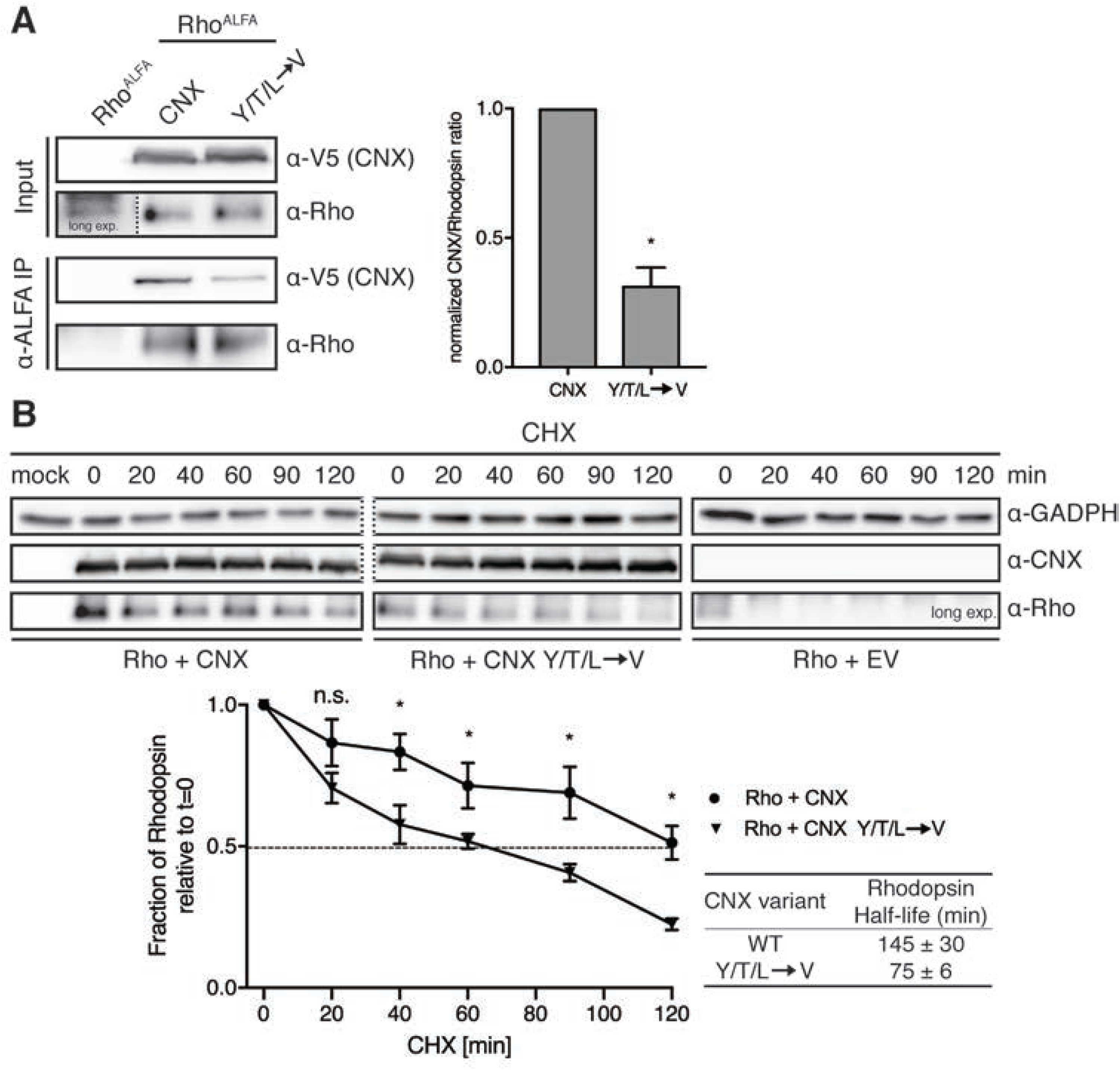
Biological functions of intramembrane client recognition by Calnexin. A Analysis of th interaction of the binding deficient CNX YTL TMD variant to Rhodopsin. CNX knockout HEK293T cells were transiently transfected with either N-terminal ALFA-tagged Rhodopsin alone or in combination with the indicated V5-tagged CNX constructs. In the absence of any CNX, Rho is hardly detectable (long exposure shown). Cell lysates and ALFA-immunoprecipitates were analyzed for Rhodopsin and co-immunoprecipitating CNX. One representative blot is shown on the left, the quantification on the right (mean ± SEM, N = 4, *P value < 0.05, two-tailed Student’s t tests). B Analysis of Rhodopsin degradation and its dependency on CNX. CNX knockout HEK293T cells that were transfected with the indicated constructs were incubated with cycloheximide and lysates were collected at different timepoints. CNX WT and CNX YTL immunoblots are from the same blot (dashed line). GADPH was used as a loading control. Quantifications are shown below representative immunoblots (mean ± SEM, N ≥ 8, *P value < 0.05, two-tailed Student’s t tests).

## Discussion

Focusing on CNX, one of the key chaperones of the ER, this study significantly advances our understanding of the molecular mechanisms that underlie membrane protein chaperoning. We show that CNX recognizes membrane protein clients in the lipid bilayer, even if non-glycosylated, which reconciles a long debate about this chaperone. Furthermore, we identify chaperone- and substrate-intrinsic motifs that allow client recognition in the membrane, which to our knowledge is the first systematic bilateral analysis on this important cell biological question. And lastly, our data show that a large variety of CNX clients are non-glycosylated and depend on CNX for their physiological levels, thus providing a new perspective on the client repertoire of this important chaperone.

Amongst the non-glycosylated membrane protein clients of CNX we find roughly equal numbers of single-pass but also multipass TM proteins. It has been suggested that the TM helix of CNX may act as a placeholder until assembly of multipass TM proteins in the lipid bilayer is complete (Cannon & Cresswell, 2001). This is consistent with our findings that CNX binds to several multipass membrane proteins. It is also consistent with our data that CNX has a preference for intra-membrane mutants of TM proteins, where helix assembly in the bilayer may fail. The large number of non-glycosylated single-pass TM protein clients of CNX we identify, however, suggests additional functions, in particular since not all of those are (known to) be part of multiprotein complexes. A likely additional function is to link substrate recognition in the membrane to ER lumenal chaperone action: whenever CNX binds to TM helices of clients in the membrane, its ER lumenal domain can recruit further folding factors like the PDI ERp57 (Oliver *et al*, 1997) or the PPIase CypB (Kozlov *et al*, 2010a). Our study suggests that this recruitment is not necessarily dependent on client glycosylation, which significantly extends the substrate repertoire of the CNX-centered folding machinery. It is noteworthy that ∼50% (1400 out of 2790 proteins) of all human multipass membrane proteins do not contain any domains outside the membrane (defined as continuous stretchers longer than 100 amino acids, TM domains predicted with Phobius (Käll *et al*, 2004) using the UniProt databank (UniProtConsortium, 2021)). In contrast, among the human proteins with only one TM domain, 90% do contain domains outside the membrane. This highlights the need for intramembrane chaperoning for multipass membrane proteins - as well as the need of recruiting folding machineries for soluble domains to single pass membrane proteins.

For CNX itself, we find a surprisingly simple principle of client engagement in the membrane: its single TMD can apparently directly bind TM clients, crucially involving a YxxTxxL motif in the CNX TM region (with x being any amino acid, although we did not analyze this further) (Fig 7). The amino acids in this motif point towards the same face of the CNX TMD, and our data show that the position of the motif within the membrane is important for client binding. It should be noted that CNX can also form ERp29-mediated dimers (Nakao *et al*, 2021), which may affect intra-membrane client binding. Our data cannot completely rule out that further factors are involved in CNX client binding, as CNX has a large repertoire of interacting proteins in the ER that e.g. involve the membrane protein chaperone EMC (Christianson *et al*, 2011). This may suggest that CNX can also act synergistically with other TM protein chaperones in the ER, although together our data favour the interpretation of direct client engagement by the CNX TMD. We thus performed a bioinformatic analysis on the occurrence of the YxxT motif in other membrane proteins in yeast and humans (leaving out the very common L residue for this analysis) of the ER and inner nuclear membrane. Of note, similar motifs are found in the TMDs of yeast Spf1, Asi2, Hrd1 and Dfm1 (Appendix Fig S7A), factors that are all involved in protein quality control and may recognize clients in the lipid bilayer (McKenna *et al*, 2020; Natarajan *et al*, 2020; Neal *et al*, 2018; Sato *et al*, 2009). In humans, among others, this motif is found in Bap31, RHBL4 (Appendix Fig S7B), which are also known to be involved in membrane protein quality control (Fleig *et al*, 2012; Geiger *et al*, 2011). This may suggest a more general role of a YxxT or similar motif in this process – and importantly now allows to test this experimentally.

**Fig 7.**
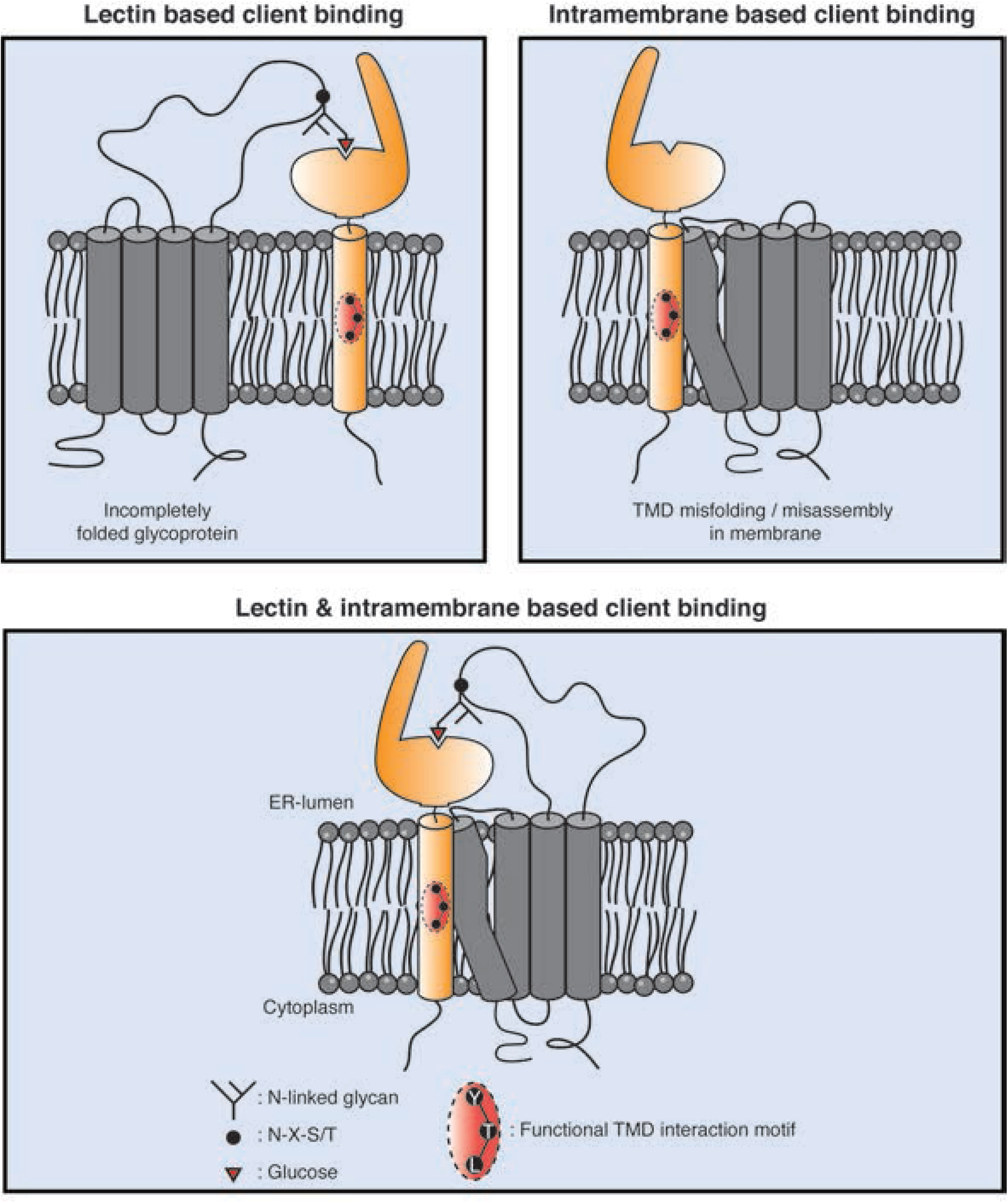
Intra-membrane client recognition potentiates the chaperone functions of Calnexin. A model for the recognition and quality control of membrane proteins by CNX. Calnexin can recognize clients via mono-glucosylated glycans (lectin-based binding mode). Additionally, membrane proteins containing a misfolded/mis-assembled TMD are recognized and bound by calnexin via a substrate interaction motif located in its TMD (intramembrane-based client binding). For glycosylated membrane proteins, both modes can work synergistically to chaperone clients.

The identification of a client recognition motif within the CNX TM region allowed us to assess the biological impact of lectin-*versus* TM-based binding of CNX. We find that for the single-pass client ConMem, binding can occur via the CNX TMD or, if ConMem was glycosylated, via the CNX lectin domain. Consistent with this, for Rhodopsin, a multipass glycoprotein, both recognition modes work synergistically: ablating binding in the membrane reduces Rhodopsin stability, but to lesser extent than complete CNX deletion from cells. This highlights the functional crosstalk between both binding modes, which, characteristic for chaperones, each need to be dynamic and of rather low affinity (Fig 7).

Recent work has revealed that several previously ill-characterized proteins act as membrane protein chaperones, including the EMC (Chitwood *et al*, 2018; Coelho *et al*., 2019; Jonikas *et al*, 2009; Satoh *et al*, 2015; Shurtleff *et al*, 2018), the PAT complex (Chitwood & Hegde, 2020; Meacock *et al*, 2002) and the Asi complex in yeast, which appears to control assembly of membrane proteins (Natarajan *et al*., 2020). Many of these factors may work synergistically to support membrane protein biogenesis (McGilvray *et al*, 2020). Our systematic analysis revealed TM-intrinsic preferences for client binding by CNX. The strongest binding is observed for a centrally located Arg residue. Arg, among all amino acids, has a high propensity to be protonated even within the lipid bilayer (Yoo & Cui, 2008). This may also affect its integration into the membrane, e.g. TMD tilting, involving snorkeling of the Arg residue towards the negatively charged headgroups of the lipid bilayer, and thus influence binding. For the PAT complex, polar residues within the TMD of Rhodopsin were found to be important for binding, again in particular arginine (Chitwood & Hegde, 2020). Of note, in addition to the PAT complex, CNX was among the most highly enriched interactors of early intermediates in Rhodposin biogenesis (Chitwood & Hegde, 2020). This suggests a concerted action of CNX and the PAT complex on membrane proteins in the process of their biogenesis, which our study now reveals not to be restricted to the lectin functions of CNX. Arg residues seem to be a hotspot of TM chaperone recognition, including CNX and the PAT complex. Of note, Arg-introducing mutations are often functionally detrimental for a membrane protein (Fink *et al*, 2012). Interestingly, apart from Arg residues, CNX and the PAT complex seem to have complementary binding preferences, with PAT favoring polar residues in the membrane, CNX disfavoring them, which may suggest a synergistic action on clients. For the EMC, chaperoning functions have also been proposed for polar TMDs (Coelho *et al*., 2019; Shurtleff *et al*., 2018; Tian *et al*, 2019) but also generally on first transmembrane helices to allow for their correct integration (Chitwood *et al*., 2018). It is noteworthy, that for none of these three factors, EMC (when acting as a chaperone), PAT or CNX, it is currently known how clients are released. It may be simply a competition between folding and binding, as suggested for the PAT complex (Chitwood & Hegde, 2020), but for soluble chaperones, generally energy-dependent processes regulate client release. In this light, the dual binding mode for CNX, via its TMD and lectin domains, is an interesting finding as it allows for synergistic binding of clients that show two chaperone recognition sites at the same time, yet will likely lead to reduced affinity if only one site is engaged, e.g. due to modification of glycans within the client or TMD folding/assembly.

Taken together, from functional (Conti *et al*, 2015) and structural studies (McGilvray *et al*., 2020) the picture of a complex molecular machinery supporting membrane protein biogenesis in proximity to the translocon emerges. Recent work together with this study now allow us to begin to systematically, mechanistically and structurally understand how individual membrane protein chaperones in this complex system recognize their clients and synergistically function to allow cells to correctly produce one third of their proteome.

## Materials and Methods

### DNA constructs

ConMem WT constructs, CNX and minCNX were obtained from GeneArt Gene Synthesis (ThermoFisher) in a pcDNA3.4 TOPO expression vector optimized for mammalian expression. The TMD sequence of ConMem WT was designed by multiple sequence alignment of 200, randomly selected, human single pass plasma membrane proteins (152 type I and 48 type II orientation) whose TMD sequences were obtained from the Membranome 2.0 Database using Unipro U Gene software. The human CNX sequence was obtained from UniProt (P27824) and complemented with a C-terminal epitope tag. For construction of the minCNX construct, the human CNX TMD sequence was obtained from UniProt, its TMD helix verified according (Hessa *et al*., 2007) to avoid artificially shortening of the TMD sequence and finally complemented by endogenous amino acids flanking the TMD region N- and C-terminally (Appendix Fig S2D). Human Cx32 and Rhodopsin cDNA was obtained from Origene and cloned into a pSVL vector (Amersham). C_L_-Cx32-TMD1 reporter constructs were synthesized by GeneArt Gene Synthesis (Thermo Fisher) and cloned into a pSVL vector. Individual construct components of the designed proteins and N- as well as C-terminal epitope tags were separated by (GGGS)2, (GSGS)2 or (GGGA)2 linkers. The full sequence of the CL-Cx32-TMD1 construct, which was mainly used in this study, is: MAWISLILSLLALSSGAISQAGQPKSSPSVTLFPPSSEELETNKATLVCTITDFYPGVVTVDWKVDGTPVTQGMETTQPSKQSNNKYMASSYLTLTARAWERHSSYSCQVTHE GHTVEKSLSRADSRGSGSGSGSGSG*WVSEAAVVLVMIRFIFIVSLWVRGIAT*GSGS GSGSGSQVTSS. For all other CL-CX32-TMD constructs, the TMD sequence shown in italic letters was replaced by the TMD sequence of TMD2, TMD3 or TMD4 of Cx32, which are shown in Fig 2A. Cloning into the mammalian pSVL expression vector was performed using restriction enzymes BamHI and XhoI followed by T4 DNA ligase (Promega) ligation. Introduction of epitope tags and point mutations was carried out via site-directed mutagenesis PCR using overlapping, complementary mutagenesis primers using Pfu polymerase (Promega) and subsequent DpnI (NEB) digestion. All constructs were verified by sequencing prior to use (Eurofins Genomics).

### Cell culture and transient transfections

HEK293T cell were cultivated in Dulbecco’s Modified Eagle’s Medium ((DMEM), high glucose (Sigma-Aldrich), supplemented with 10% (v/v) fetal bovine serum (Biochrom) and 1% (v/v) antibiotic-antimycotic solution (25 µg/ml amphotericin B, 10 mg/ml streptomycin, 10,000 units of penicillin (Sigma-Aldrich)) (complete DMEM) at 37°C and 5% CO_2_. 24 h prior to transfection, 250.000 HEK293T cells were seeded per p35 plate (pre-coated Corning BioCoat Poly-D-Lysine 35 mm #354467 or uncoated Nunclon multidish 6 well plates, Thermo Scientific #140685 coated with 50 µg/ml Poly-D-Lysine solution (Gibco A38904-01) per well according to manufacturers instructions) or 300.000 per p60 plate (Tissue Culture Dish 60, TPP). Transient transfection was performed using GeneCellin (Eurobio) according to the manufacturer’s protocol. For transient transfections using pcDNA (strong CMV promotor) the amount of DNA was reduced to half the amount suggested by the manufacturer.

### Cycloheximide chase assays

Cells were transfected with the indicated constructs. Inhibition of further protein biosynthesis and determination of protein half-lives was performed by the replacement of complete medium with complete medium containing 50 µg/ml Cycloheximide (Sigma-Aldrich) 24 h after transfection and the collection of lysates, as described below, at different time points. Controls were prepared in the same manner without the addition of CHX (t = 0 sample).

### Induction of ER stress

The induction of the unfolded protein response was carried out by the addition of 5 µg/ml tunicamycin (Sigma-Aldrich) in medium and incubation for 6 h prior cell lysis or the supplementation of pre-warmed medium with 10 mM DTT (Sigma-Aldrich) for one hour. Cleavage analysis of the activating transcription factor 6 (ATF6) based on immunoblotting was performed following immunoprecipitation to increase the amount of detectable protein. Determination of endogenous phosphorylation by immunoblotting (total amount and phosphorylation at Ser51 only) of the eukaryotic initiation factor 2α (eIF2α) was performed following cell lysis using lysis buffer additionally supplemented with 1 x phosphatase inhibitor mix (Serva). To assess XBP1 splicing, RNA was extracted using RNeazy Mini Kit (Qiagen) in an RNase free environment. Subsequently RT-PCR of the purified RNA was performed using OligodT20 Primer (18418020,Thermo Fisher) and SuperScript III Reverse Transcriptase (18080044, Thermo Fisher). Following amplification of the XBP1 transcript of the resulting cDNA, XBP1 splicing events were analyzed on 2% agarose gels.

### DSSO Crosslinking

Transiently transfected cells were washed twice with ice-cold PBS prior the addition of 2 mM DSSO (Thermo Fisher) in PBS per sample, diluted from 100 mM stock DSSO in anhydrous DMSO (Thermo Fisher). Crosslinking was performed on ice for 1 hour including regular agitation. Inhibition of the crosslinking reaction was achieved by washing of the samples with quenching solution (50 mM Tris-HCl pH 8.0) for 15 min. In the following, cell lysis, immunoprecipitation and sample preparation for mass spectrometry was performed as described below.

### Cell lysis

Cells were harvested 24 h after transfection (if not described otherwise). All steps were performed on ice or at 4°C using ice-cold solutions. Cells were washed using PBS and then lysed for 20 min by adding 1 ml (p60) or 500 µl (p35) lysis buffer (50 mM Tris-HCl pH 7.5, 150 mM NaCl and 1% Digitonin (Sigma-Aldrich) or 0.5% NP-40 (Sigma-Aldrich) complemented with 0.5% DOC for ConMem, Cx32, CNX and minCNX constructs. For MS experiments, lysis buffer containing 1% NP-40, 1mM MgCl_2_ and 5% Glycerol was applied to the cells. C_L_ Cx32-TMD constructs were lysed in NP-40 buffer only when conducting co-immunoprecipitation experiments otherwise Digitonin based buffer was used for all other experiments. All Digitionin and NP-40 based buffers were supplemented with cOmplete protease inhibitor cocktail (Roche) prior to lysis. The resulting lysate was centrifuged for 15 min at 15,000 x g and the supernatant was used for subsequent experiments.

### Deglycosylation experiments

Samples were digested for 1 h at 37°C with either EndoH_f_, or PNGase F (NEB) according to the manufacturer’s instructions. Negative controls were prepared in the same manner without the addition of enzymes. The digested proteins were thereafter supplemented with 5x Laemmli buffer and 2% (v/v) β-mercaptoethanol and boiled for 10 min at 95°C.

### Immunoprecipitation

Prior to co-immunoprecipitation (co-IP), 2% of cell lysate were supplemented with 5x Laemmli buffer containing 2% (v/v) β-mercaptoethanol as input samples. The remaining lysate was incubated for 2 h with 1.5 µg antibody and subsequently for 1 h with 30 µl protein A/G agarose beads (sc-2003, Santa Cruz), or for 3 h either with 25 µl of M2 anti-FLAG affinity gel (A2220, Sigma-Aldrich) or RFP-Trap Agarose (rta-10, Chromotek) or ALFA Selector ST Agaraose (N1511, NanoTag) under rotation at 4°C. Beads were thereafter washed three times with 1 ml wash buffer (50 mM Tris/HCl, pH 7.5, 150 mM NaCl and 0.5 % Digitonin (Sigma-Aldrich) or 0.5% NP-40 (Sigma-Aldrich) complemented with 0.5% DOC for ConMem, Cx32, CNX or minCNX constructs. For C_L_ Cx32-TMD constructs 0.5 % GDN (Anatrace) was used and centrifugation steps for 1 min at 8,000 x g and 4°C. Proteins were then eluted by addition of 22 µl (RFP-Trap) or 33 µl of 2x Laemmli supplemented with β-mercaptoethanol and subsequent incubation at 95°C for 10 min, or 50°C for all samples containing ConMem and 37°C for 30 min for all samples containing full-length Cx32.

### SDS-PAGE and immunoblotting

Proteins were separated by SDS-polyacrylamide gel electrophoresis (PAGE) in 10% to 14% SDS-PAGE gels. The polyacrylamide gels were then either imaged directly using a Typhoon 9200 Variable Mode gel scanner (GE Healthcare) (Cy5 filter setting, excitation 633 nm, emission 670 nm, bandwidth 30 nm or GFP excitation 526 nm, emission 532 nm, short pass) or subsequently used for western blots. Proteins were blotted overnight at 4°C onto polyvinylidene difluoride (PVDF) membranes (Biorad) and subsequently, the membranes were blocked for 6 h at RT (or overnight at 4°C) with Tris-buffered saline supplemented with skim milk powder and Tween-20 (M-TBST; 25mM Tris/HCl, pH 7.5. 150 mM NaCl, 5% (w/v) skim milk powder, 0.05% (v/v) Tween-20). Primary antibodies in M-TBST were applied at 4°C overnight. After washing (1 x 5 min TBS, 2 x 5 min TBST, 3 x 5 min TBS), the blots were decorated for 1 h at RT with the respective HRP-conjugated secondary antibody diluted in M-TBST (at 10-fold lower dilution than the respective primary antibody). Blots were imaged using Amersham ECL prime solution (GE Life Sciences) and a Fusion Pulse 6 imager (Vilber Lourmat). Quantifications were conducted using Bio-1D software (Vilber Lourmat).

### Antibodies

For western blots, the following primary antibodies were used at the dilutions listed: rabbit monoclonal anti-EMC4 (ab 184544, Abcam) at 1:5,000; mouse monoclonal (M2) anti-FLAG (F1804, Sigma-Aldrich) at 1:1,000; mouse monoclonal (16B12) anti-HA (901513, Biolegend) at 1:500; mouse monoclonal (C8.B6) anti-calnexin (MAB3126, Chemicon) at 1:1,000; monoclonal rat (W17077C) anti-calnexin (699402, Biolegend) at 1:1,000; mouse monoclonal (B-6) anti-Hsc70 (sc-7298, Santa Cruz) at 1:1,000; polyclonal rabbit anti-connexin32 (C3595, Sigma-Aldrich) at 1:500; mouse monoclonal (6G6) anti-RFP (chromotek) at 1:1,000; polyclonal goat anti-mouse lambda (1060-01, Southern Biotech) at 1:250; monoclonal mouse anti-Rhodopsin (MA1-722, Invitrogen) at 1:250; mouse monoclonal anti-ATF6 (ab122897, Abcam) at 1:500; polyclonal rabbit anti-(Ser51)elF2alpha (9721, Cell Signaling) at 1:500; polyclonal rabbit anti-eIF2alpha (9722, Cell Signaling) at 1:1000; polyclonal rabbit anti-BiP (C50B12, Cell Signaling) at 1:500; polyclonal rabbit anti-SPCS2 (14872-1-AP, Proteintech) at 1:1000; polyclonal rabbit anti-ATP6AP2 (HPA003156, Sigma-Aldrich) at 1:250. The following HRP-conjugated secondary antibodies or proteins were used for development of western blots at 1:5000 – 1:10000: mouse IgGκ-binding protein (m-IgGκ BP-HRP) (sc-516102, Santa Cruz); mouse monoclonal anti-rabbit IgG-HRP (sc-2357, Santa Cruz); mouse monoclonal anti-goat IgG-HRP (sc-2354, Santa Cruz); goat polyclonal anti-rat IgG (Poly4054, Biolegend). For IP, the following antibodies were employed: polyclonal goat anti-mouse lambda (1060-01, Southern Biotech); mouse monoclonal (C8.B6) anti-calnexin (MAB3126, Chemicon); polyclonal rabbit anti-HA.11 (Poly9023, Biolegend).

### Immunofluorescence

#### Seeding and transfection

36 µl DMEM containing 3.6 µg DNA were mixed with 1.2 µl TorpedoDNA transfection reagent (ibidi) and incubated for 15 min at RT. 200 µl of COS-7 cell suspension with 4 x 10^5^ cells/ml were added and mixed gently. 30 µl of the resulting suspension were applied per inlet of a µ-Slide IV 0.4 (ibidi) and the µ-Slide was incubated for 3 h at 37°C, 5% CO_2_. According to manufactureŕs instructions, medium was replaced 3 h after seeding. Therefore, 60 µl complete DMEM were added per reservoir, and the µ-Slides were incubated for additional 21 h at 37°C, 5% CO_2_.

#### Staining

For live cell imaging, medium was removed from µ-Slide reservoirs and each channel washed twice with 100 µl 37°C PBS. Subsequently, samples were imaged. For fixation, all liquid was removed from all reservoirs and channels and 60 µl glyoxal fixation solution (20% EtOH, 7.825% glyoxal, 0.75% acetic acid) (Richter *et al*, 2018) were added. Samples were then incubated for 30 min on ice and thereafter further 30 min at RT. The reaction was quenched by aspirating the fixation solution and adding 60 µl of 100 mM NH_4_Cl (Sigma-Aldrich) and subsequent incubation at RT for 20 min. Samples were then washed twice for 5 min with 100 µl 4°C PBS. 60 µl of blocking solution (2.5% BSA (Sigma-Aldrich), 0.1% Triton X-100 in PBS) were added and samples were then incubated for 5 min at RT following three further washing steps at RT. For FLAG-tagged constructs, 30 µl of anti-FLAG mouse monoclonal antibody (F1804, Sigma-Aldrich) at 1:500 dilution in blocking solution were added and incubated for 2 h at RT and subsquenctly washed three times for 5 min with 100 µl PBS. As an ER-marker, 30 µl of anti-PDI antibody conjugated to Alexa Fluor® 488 (#5051, Cell Signaling Technology) at 1:50 dilution in blocking solution were added and incubated for 1 h at RT in the dark. For FLAG-tagged constructs, donkey polyclonal anti-mouse IgG antibody conjugated with TexasRed (PA1-28626, Thermo-Fisher) at 1:300 dilution and for ALFA tagged CNX constructs ALFA-FluoTag-X2 nanobodies (NanoTag Biotechnologies) at 1:250 dilution were added to the PDI-staining solution. The antibody solution was washed out using 100 µl PBS and the samples were thereafter washed three times. All liquid was removed and 25 µl of DAPI solution (Sigma-Aldrich) (0.01% in PBS) were added to stain Nuclei and incubated for 2 min. The samples were then washed three times with PBS, the liquid was aspirated, and mounting medium (ibidi) was added to cover the inlets of the slides.

#### Microscopy

Imaging was performed on a DMi8 CS Bino inverted widefield fluorescence microscope (Leica) using a 100 x (NA = 1.4) or 63 x (NA = 1.4) oil immersion objective. The employed dichroic filters were chosen to image Alexa 488 and sfGFP (GFP channel; excitation/bandpass: 470/40 nm; emission/bandpass: 525/50 nm), mScarlet-I, Sec61mCherry and TexasRed (TXR channel; excitation/bandpass: 560/40 nm; emission/bandpass: 630/75 nm), or DAPI (excitation/bandpass: 350/50 nm; emission/bandpass: 460/50 nm). For image analysis and processing, the LAS X (Leica) analysis software and ImageJ (NIH) where used. Adjustments of acquired images were restricted to homogenous changes in brightness and contrast over the whole image.

### Quantification and statistics

Immunoblots were quantified using the Bio-1D software (Vilber Lourmat). Binding of CNX to full-length Cx32 WT and mutants, individual Cx32-TMD domains, Rhodopsin or ConMem TMD helices was calculated as the ratio of intensities of the co-immunoprecipitated protein (CNX) to the overall intensity of immunoprecipitated protein. In case of ConMem the overall intensity of immunoprecipitated protein was determined as the sum of both glycosylated and nonglycosylated species – beginning at the molecular weight of HA-tagged nonglycosylated ConMem until the upper protein smear represented by the complex glycosylated species. In case of ConMem N_1_Q, the area around the nonglycosylated species was included into quantification. Subsequently, to facilitate comparability of individual experiments, formed ratios were each normalized to the ConMem N_1_Q (ConMem lacking the lumenal glycosylation site) dataset of each experiment or CNX when examining binding to the binding deficient variants. The decay following CHX treatment was calculated by normalizing the quantified intensity at each time point to the intensity of the control sample (t = 0 min) not treated with CHX. Determination of half-life was performed by logarithmic linearization of the CHX decays and determining the point of intersection of this line to ln (0.5). Statistical analyses were performed using Prism (GraphPad Software). Where indicated, data were analyzed with two-tailed, unpaired Student’s t tests and differences were considered to be statistically significant when P < 0.05.

### Sequence analysis and structural modeling

TMD regions of Cx32 and ConMem WT as well as mutant variants were annotated by using the ΔG prediction server v1.0 (Stockholm University). The same tool was used to predict the ΔGapp for helix insertions using the full protein scan mode. Structural analysis of and identification of lysine residues as potential DSSO crosslinking targets within the lumenal domain of CNX was performed using PyMol and mammalian (Canis lupus familiaris, (Schrag *et al*, 2001)) CNX as template. Helical wheel projection of the CNX TMD sequence was performed using the web based NetWheels application (http://lbqp.unb.br/NetWheels/).

### Bioinformatic analyses

Reference proteomes of Homo sapiens (UP000005640) and Saccharomyces cerevisiae (UP00000231) were downloaded from the UniProt database (Consortium, 2021). Subcellular locations were assigned to the proteins according to their UniProt annotation. Only proteins that have been found to be located in the endoplasmic reticulum or nucleus inner membrane were taken into the analysis. Sequence positions of transmembrane (TM) regions were predicted using Phobius (Käll *et al*., 2004). The YxxT motifs were detected in the TM sequences by matching the corresponding regular expression (script in python). The first one and last three residues of TM regions were excluded from the search for the motif.

### Gene knockouts

CNX-deficient cells were generated using the CRISPR/Cas9 system. The vector pSpCas9(BB)-2A-Puro (PX459) V2.0 was provided from Feng Zhang (plasmid 62988; Addgene, Cambridge MA) (Ran *et al*, 2013). The design of guide RNA was performed using the German Cancer Research Center (DKFZ) E-CRISP design tool. In order to prevent the occurrence of off-target effects two different guide RNA sequences for the CNX target gene were designed (gRNA #1: 5‘-CACCGCTTGGAACTGCTATTGTTG-3’; gRNA #2: 5‘- CACCGTGGTTGCTGTGTATGTTAC-3’) and subsequently cloned into PX459 according to published protocols (Ran *et al*., 2013). In the following, HEK293T cells were transfected using GeneCellin (Eurobio) according to the manufacturer’s protocol and cultured for two days. Selection of cells was carried through to the addition of 1.5 µg/mL puromycin for 72 hours. Subsequently, single colonies were isolated and CNX protein levels were determined by immunoblotting. To ensure a complete knockout of the target gene and loss of the respective protein this process was repeated during several passages. Furthermore, following the amplification of the genomic CRISPR/Cas9 target area by the usage of specific primers, the obtained amplicons were sequenced and the successful CNX knockout verified by NGS CRISPR amplicon sequencing (CCIB DNA Core, Massachusetts General Hospital, Boston, MA).

### MS sample preparation and measurement of CNX enrichment

HEK293T CNX KO cells were seeded in P100 plates and cultivated for 24 h prior transfection with 10 µg of EV or ALFA-tagged CNX variants with a total of three replicates for each construct. DSSO crosslinking and cell lysis was performed as described above using NP-40 lysis buffer additionally supplemented with 1 mM MgCl_2_ and 5% Glycerol. Immunoprecipitations were performed in the same buffer using anti-ALFA Selector ST agarose beads as described above followed by two additional washing steps in detergent free buffer. During further proceedings, immunoprecipitated proteins destined for MS analysis were digested and eluted from beads prior to desalting and purification of the samples as otherwise described (Keilhauer *et al*, 2015). Nanoflow liquid chromatography-mass spectrometry (MS)/MS analyses were performed using a combination of an UltiMate 3000 Nano HPLC system (Thermo Fisher) together with an Orbitrap Fusion mass spectrometer (Thermo Fisher). Peptides were first loaded on an Acclaim C18 PepMap100 75 µm ID × 2 cm trap column together with 0.1% FA before transfer to an Aurora reversed phase UHPLC analytical column, 75 µm ID × 25 cm, 120 Å pore size, (Ionopticks). Columns were constantly heated at 40 °C. Subsequent separation was performed using a first gradient ranging from 5 to 22% acetonitrile in 0.1% FA for 105 min followed by a second gradient ranging from 22 to 32% acetonitrile in 0.1% FA for 10 min at an overall flow rate of 400 nl/min. Peptides were ionized via electrospray ionization. Orbitrap Fusion was carried out in a top speed data dependent mode using a cycle time of 3 s. Full scan (MS^1^) acquisition (scan range of 300 – 1500 m/z) was performed in the orbitrap at a defined resolution of 120,000 as well as with an automatic gain control (AGC) ion target value of 2e5 whereby dynamic exclusion was set to 60 s. For fragmentation, precursors with a charge state of 2-7 and a minimum intensity of 5e3 were selected and isolated in the quadrupole using a window of 1.6 m/z. Subsequent fragment generation was achieved using higher-energy collisional dissociation (HCD, collision energy: 30%). The MS^2^ AGC was adjusted to 1e4 and 50 ms were selected as the maximum injection time for the ion trap (with inject ions for all available parallelized time enabled). Scanning of fragments was performed by applying the rapid scan rate.

### MS search and bioinformatics for CNX enrichment

Thermo raw files were analyzed with MaxQuant software (version 1.6.17.0) with most default settings and a protein database containing human sequences (downloaded July 2019 from Uniprot, taxonomy ID: 9606, 74349 entries). The ALFA tagged version of Calnexin was not implemented. The following parameter settings were used: PSM and protein FDR 1%; enzyme specificity trypsin/P; minimal peptide length: 7; variable modifications: methionine oxidation, N-terminal acetylation; fixed modification: carbamidomethylation. The minimal number of unique peptides for protein identification was set to 2. For label-free protein quantification, the MaxLFQ algorithm was used as part of the MaxQuant environment: (LFQ) minimum ratio count: 2; peptides for quantification: unique. Statistical analysis was performed in Perseus (version 1.6.14.0). Proteins identified only by site, reverse hits or potential contaminants were removed. LFQ intensities were log2 transformed and data were then filtered for at least two valid values in at least one group (wt). Then, missing values were imputed from normal distribution (width: 0.3, down shift: 1.8 standard deviations, mode: over whole matrix). The replicate groups were compared via a two-sided, two-sample Student’s t test (S0 = 0, permutation-based FDR method with FDR = 0.05 and 250 randomizations). Enrichment values and corresponding −log10 P values were plotted. Cut offs in the volcano plot were set to 2-fold enrichment and a p-value of 0.05.

### MS sample preparation and measurement parameters: proteome

HEK293T CNX KO cells were seeded in P100 plates and transfected with empty vector or reconstituted with vector coding for CNX WT. Four plates of either condition were used for the four independent replicates.

Lysis was performed two days after transfection by removing the supernatant and adding 300 µl of lysis buffer (as described above) to the cells followed by 10 min incubation on ice. Cells were removed from plates by scraping and precleared by centrifugation at 13000 rpm for 10 min at 4°C. Protein concentration was determined using a BCA assay (Life Technologies) and 300 µg of each sample was precipitated according to (Wessel & Flügge, 1984) and proteins afterwards resuspended in 8M urea, 0.1 M Tris/HCl pH 8.5 for denaturation. Samples were loaded on molecular cut-off spin columns (Microcon 30 kDa, Merck Millipore) and subsequently reduced, alkylated and digested with trypsin on the filters following the FASP protocol described by (Wiśniewski *et al*, 2009). The obtained peptide mixtures were lyophilized, desalted on 50 mg SepPac C18 columns (Waters) and afterwards fractionated on self-made SCX stage tips (3 disks of Cation extraction material, Empore, in 200 µl pipette tips). Peptides bound to the SCX material were step wise eluted with five different concentrations of AcONH4 (20 mM, 50 mM, 100 mM, 250 mM and 500 mM), 0.1 % TFA, 15 % acetonitrile. Overall 40 samples (5 fractions for each of the two conditions and four replicates) were further evaporated in a Speed Vac, desalted on self-made C18 stage tips (2 layers of C18 disks, Empore) and after final evaporation resuspended in 25 µl 0.1% formic acid (FA) and filtered through equlibrated 0.2 μm Millipore filters before MS measurement.

Nanoflow liquid chromatography-mass spectrometry (MS)/MS analysis for the proteome analysis was performed on a TimsTOF Pro mass spectrometer equipped with an CaptiveSpray nano-electrospray ion source (Bruker Daltonics) coupled to an UltiMate 3000 Nano HPLC system (Thermo Fisher). The column setup for peptide separation was as described above. Gradient length on the TimsTOF Pro was 73 mins, while acetonitrile in 0.1 % FA was step wise increased from 5 to 28% in 60 mins and from 28 to 40% in 13 mins, followed by a washing and equilibration step of the column.

The timsTOF Pro was operated in PASEF mode (Meier *et al*, 2015; Meier *et al*, 2018).

Mass spectra for MS and MS/MS scans were recorded between 100 and 1700 m/z. Ion mobility resolution was set to 0.85–1.40 V·s/cm over a ramp time of 100 ms. Data-dependent acquisition was performed using 10 PASEF MS/MS scans per cycle with a near 100% duty cycle. A polygon filter was applied in the m/z and ion mobility space to exclude low m/z, singly charged ions from PASEF precursor selection. An active exclusion time of 0.4 min was applied to precursors that reached 20,000 intensity units. Collisional energy was ramped stepwise as a function of ion mobility.

### MS search and bioinformatics for proteome analysis

TimsTOF raw files were loaded into MaxQuant (software version 2.0.3.0) and the default settings for TimsTOF files were applied except that the TOF MS/MS match tolerance was set to 0.05 Da. For the Andromeda search the human protein database downloaded from Uniprot in September 2021 was used. Other search parameters were set as described for the CNX enrichment dataset and also statistical analysis, performed in Perseus (version 1.6.15.0) followed the same steps. Data filtration was chosen for at least three valid values in at least one group, followed by imputation of missing values and comparison of replicate groups of CNX knock out cells versus CNX reconstituted cells via a two-sided, two-sample Student’s t test (S0 = 0, permutation-based FDR method with FDR = 0.05 and 250 randomizations). Cut offs were set for the p-value < 0.05 and a difference in fold change of 20%.

### Molecular dynamics / computational modeling

#### MD Simulations

All MD simulations were performed with the GPU accelerated version of PMEMD (Salomon-Ferrer *et al*, 2013), part of the AMBER18 package (www.ambermd.org). Amino acids were described with ff14SB (Maier *et al*, 2015), lipids with LIPID17, a successor of LIPID14 (Dickson *et al*, 2014) and water molecules with TIP3P (Jorgensen *et al*, 1983). The sequences of the TM helices used for the simulations are given below. All simulations were prepared with the membrane builder (Wu *et al*, 2014) of CHARMM-GUI (Jo *et al*, 2008), utilizing the AMBER-FF compatibility (Lee *et al*, 2020). The simulation boxes consisted of approx. 200 POPC molecules, 13500 water molecules with box dimensions (after equilibration and pressurization) of 84Åx84Åx95Å. Simulations were performed in a 0.15M KCl solution. Prior to sampling, the systems were treated with the equilibration protocol, suggested by CHARMM-GUI (see Table below).

|  | Steps | Time Step | K Protein | K Lipid | K Dihed. | Temperature | Pressure |
| --- | --- | --- | --- | --- | --- | --- | --- |
| min | 2000 | minimization | 10.0 | 2.5 | 250 | --- | --- |
| Eq1 | 125000 | 1fs | 10.0 | 2.5 | 100 | 303.15K | NVT |
| Eq2 | 125000 | 1fs | 5.0 | 2.5 | 50 | 303.15K | NVT |
| Eq3 | 125000 | 1fs | 2.5 | 1.0 | 50 | 303.15K | 1 bar |
| Eq4 | 250000 | 2fs | 1.0 | 0.5 | 50 | 303.15K | 1bar |
| Eq5 | 250000 | 2fs | 0.5 | 0.1 | 25 | 303.15K | 1bar |
| Eq6 | 250000 | 2fs | 0.1 | 0.0 | 0 | 303.15K | 1bar |

Overview over the seven equilibration steps performed for all simulations. The initial minimization is followed by six equilibration simulations with decreasing force constants (K) on positional restraints of amino acids (Protein, all atoms), positional restraints on lipid headgroups (Lipid, phosphorus atom) as well as restraints on the lipid dihedrals (Dihed.).

In all cases, periodic boundary conditions were used and the temperature was set to 303.15K using the Langevin thermostat (Goga *et al*, 2012) with a friction coefficient of 1.0 ps^-1^. To sample NpT ensembles, the Berendsen manostat (Berendsen *et al*, 1984) was utillized and set to 1.0 bar with a relaxation time of 1ps. All non-bonded interactions were calculated directly up to a distance of 9Å, after which electrostatic interactions were treated with the particle mesh Ewald method (Darden *et al*, 1993), while long range van-der-Waals interactions were estimated by a dispersion correction model. To achieve time-steps of 4fs, the Shake algorithm (Anderson, 1983) as well as hydrogen mass repartitioning (Hopkins *et al*, 2015) were used. Prior to building the dimeric simulation systems, the CNX, Cx32 and ConMem helices were simulated in isolation for 1µs in a POPC bilayer in order to equilibrate the structure and to predict their orientation in the membrane. All simulation snapshots were rendered using VMD (Humphrey *et al*, 1996).

### CNX-ConMem dimerization simulations

Initially, five simulations of one CNX helix and one ConMem helix in the same simulation box, were performed. Each of the five simulations was started with different relative orientations of the two helices which were placed at random positions. After preparing the simulation systems using CHARMM-GUI, the systems were equilibrated using the aforementioned protocol. Subsequently, unrestricted sampling simulations of at least 2.5µs were performed.

Only one of the starting structures led to the formation of a stable complex. In order to inprove statistics, two more simulations were started from this initial conformation. Altogether, two successful dimerization events have been observed in these simulations. All simulations have been sampled for at least 4µs.

To obtain binding statistics on the amino-acid level that are more generally valid, we also performed simulations with CNX and two ConMem mutants: V13P and V13R. Several at least 4µs long simulations were performed for both mutants, however, only two resulted in the formation of CNX-ConMem interactions along the entire TMD: One simulation featuring V13R and one with the V13P mutant. Of interest: While the V13P mutant binds to CNX in a fashion very similar to WT ConMem, the V13R mutant uses an alternative interface to bind to CNX. CNX, on the other hand, binds all ConMem variants utilizing basically the same binding interface. Altogether, 22 µs time-scale simulations of this system have been performed. However, for analysis we only selected the 4 simulations that exhibited broad TMD-TMD interactions between CNX and ConMem. Data analysis was performed on 4µs long trajectories in each case.

### CNX-Cx32 dimerization simulations

For this system five, at least 4µs long simulations were performed. The simulation set-up was identical to the one used for the CNX-ConMem system. In four out of the five cases, we were able to sample at least transient interactions between the helices, involving the entire TMD of CNX. These four simulations were then selected for data analysis on 4µs of the sampled trajectories.

<u>CNX SEQUENCE in Simulations:</u> [ACE GLN MET ILE GLU ALA ALA GLU GLU ARG PRO TRP LEU TRP VAL VAL TYR ILE LEU THR VAL ALA LEU PRO VAL PHE LEU VAL ILE LEU PHE CYS CYS SER GLY LYS LYS GLN THR SER GLY MET GLU TYR NME]

<u>Cx32-TMD1 SEQUENCE in Simulations:</u> [ACE SER GLY SER GLY SER GLY SER GLY SER GLY THR ALA ILE GLY ARG VAL TRP LEU SER VAL ILE PHE ILE PHE ARG ILE MET VAL LEU VAL VAL ALA ALA GLU SER VAL TRP GLY SER GLY SER GLY SER GLY SER GLY SER GLY NME]

<u>ConMem (WT) SEQUENCE in Simulations:</u> [ACE GLY GLY GLY GLY SER PRO ALA ALA ILE LEU VAL ILE VAL VAL VAL VAL VAL VAL PHE ILE ILE ILE VAL VAL VAL VAL VAL PHE ILE ILE LYS GLY GLY GLY GLY SER NME]

## Acknowledgments

We are grateful to Carolin Klose (TUM), Bastian Bräuning (MPI of Biochemistry) and Marius Lemberg (University of Cologne) for helpful comments on the manuscript. The Sec61 mCherry contruct was a kind gift of Gia Voeltz, University of Boulder, Colorado. SAS gratefully acknowledges funding of the European Research Council (ERC) and the European Union’s Horizon 2020 research and innovation program (grant agreement no. 725085, CHEMMINE, ERC consolidator grant). MJF gratefully acknowledges funding by a German Research Foundation (DFG) research grant, number FE1581/1-2.

## Author Contributions

NB and MJF conceived the study and designed experiments. Mass spectrometry experiments were performed by NBa. NB, KMB and JPLC performed all other experiments. The experimental data was analyzed by NB, KMB, JPLC, NBa, SAS and MJF. Bioinformatic analyses were performed by MP and DF. MD simulations and their analysis was performed by MH and MZ. The paper was written by NB and MJF with input from all other authors.

## Appendix Information

### Supplemental Figures and Tables

**Appendix Fig S1.**
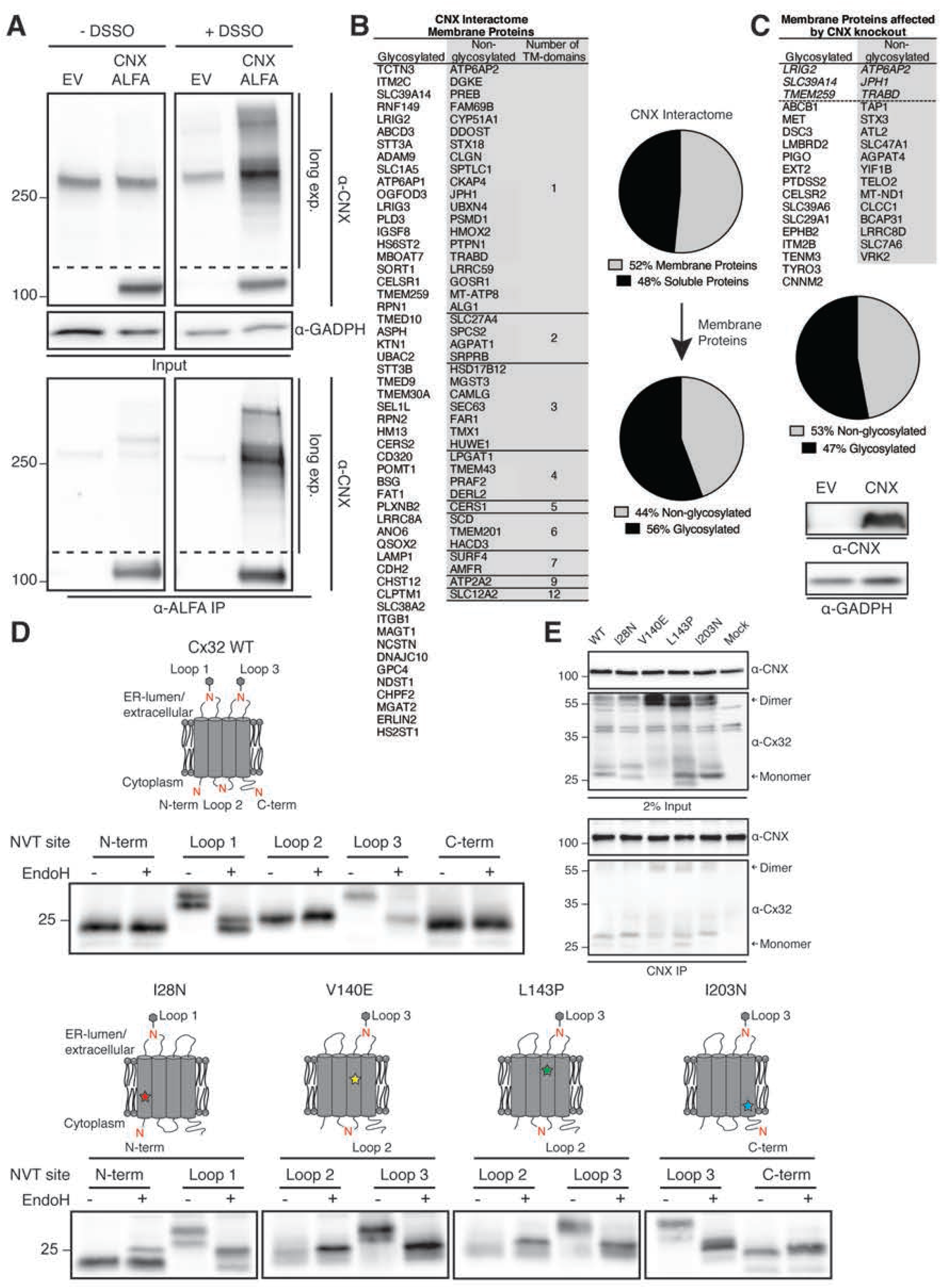
CNX client interactions via mass spectrometry and Cx32 as a model protein. **(A)** Immunoblot analysis following *in situ* DSSO crosslinking in CNX deficient cells transiently transfected with empty vector (EV) or CNX containing a C-terminal ALFA-tag. Cell lysates and ALFA-tag immunoprecipitates were immunoblotted against CNX to detect co-immunoprecipitating DSSO-induced crosslinks. GAPDH was used as a loading control. **(B)** List of identified CNX interaction partners derived from LC-MS/MS analysis of ALFA-tagged CNX, immunoprecipitated in 1% NP-40 buffer from transfected CNX deficient HEK293T cells after DSSO crosslinking and compared to empty vector control co-IPs. Significantly enriched CNX interacting proteins (-log_10_ = (P-value) > 1.3 (P<0.05) and ratio CNX WT/EV (log_2_) > 1) were used, discarding mitochondrial proteins according to the GO-term annotation “*mitochondrion*” (GO:0005739). The excluded hits were additionally screened for correctness of this annotation according to their Uniport entries. Subsequently, the resulting interactome was divided into soluble and membrane proteins through identification of membrane proteins via the GO-term annotation “*integral to membrane”* (GO:0016021). Addtionally, interaction partners, which possess TMDs but are not listed under the GO-term annotation “*integral to membrane*” were added manually to this category by Phobius analysis (Käll *et al*, 2004), a membrane topology prediction program. Listed are the identified membrane proteins and their corresponding Uniprot gene names interacting with CNX divided into glycosylated and non-glycosylated membrane proteins according to their Uniprot entries. In addition, the number of TM helices for non-glycosylated membrane proteins based on their Uniprot entries is given. **(C)** List of identified membrane proteins and their corresponding Uniprot gene names affected by CNX knockout on a proteome-wide scale derived from LC-MS/MS analysis of CNX deficient HEK293T cells and compared to control cells re-constituted with CNX WT. The identified proteome of downregulated proteins (-log p-value > 1.3 and ratio EV / CNX WT > 0,27) was screened for membrane proteins according to the GO-term annotation “*integral to membrane*” (GO:0016021), discarding mitochondrial proteins according to the GO-term annotation “*mitochondrion*” (GO:0005739). The excluded hits were additionally screened for correctness of this annotation according to their Uniprot entries. Integral membrane proteins not listed under the aforementioned GO term annotation but possessing a TM domain were manually added to the pool of IMPs affected by CNX knockout after additional analysis with Phobius (Käll *et al*., 2004). Subsequently, the individual hits were divided into glycosylated and non-glycosylated membrane proteins according to their Uniprot entries. Previously identified CNX interactors (**B**) are shown in italic and separated by a dashed line. A representative blot of cell lysate blotted against CNX and GADPH from one of the four individual replicates analyzed in MS is shown below. **(D)** EndoH digest of Cx32 WT and its mutants used in this study to verify their proper orientation within the membrane. Schematics of mutations studied, membrane integration, and reporter site glycosylation are shown (N: NVT reporter sites for glycosylation, which were individually introduced. Only the most N-terminal site (located in the cytoplasm) is endogenously present in Cx32). **(E)** HEK293T cells were transfected with Cx32 WT or the indicated mutants. Immunoprecipitations of endogenous CNX reveal interactions with monomeric and dimeric Cx32 species.

**Appendix Fig S2.**
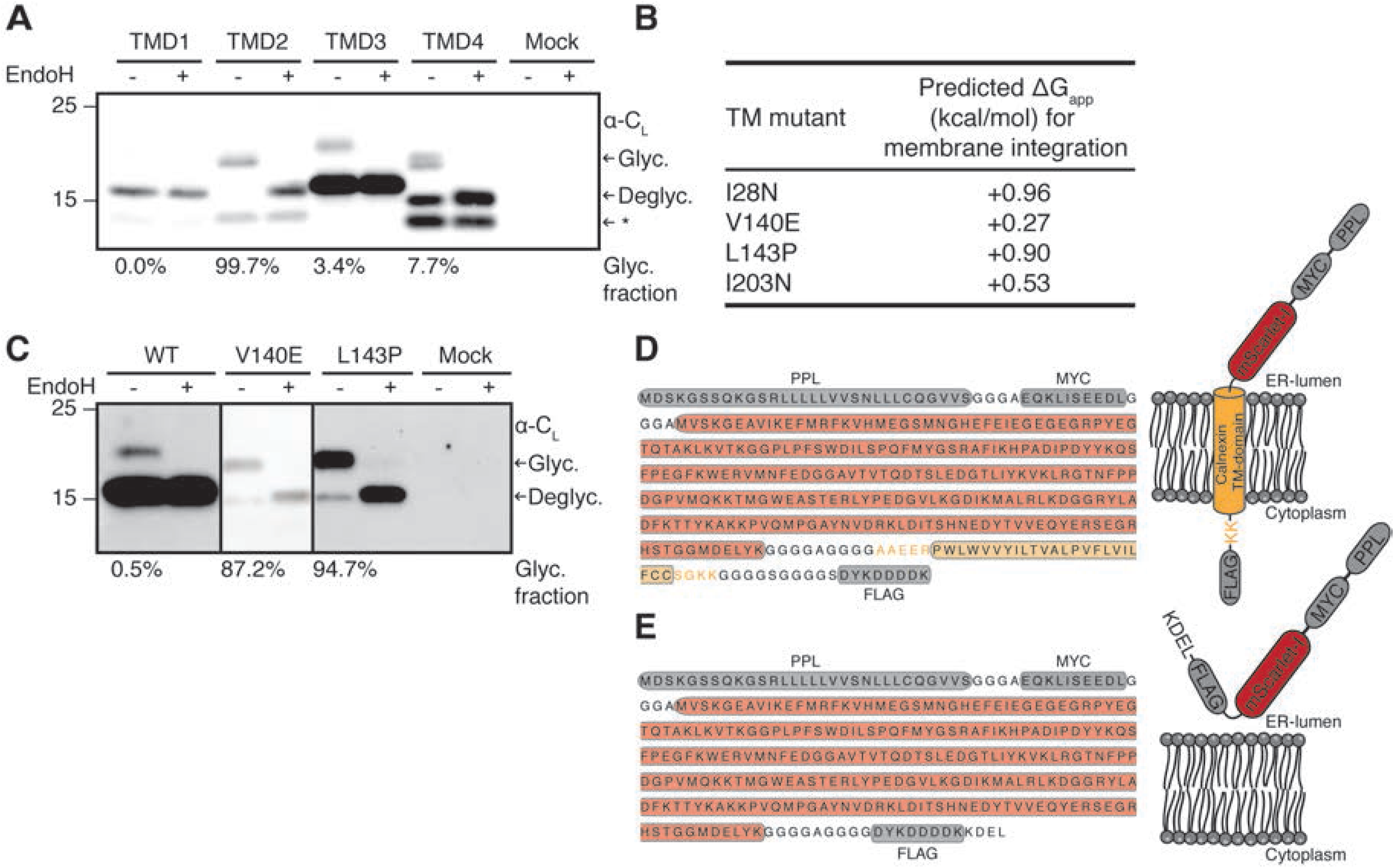
Interaction of client and CNX TMDs. **(A)** EndoH digest of the four C_L_-TMD reporters to assess their integration and orientation in the membrane. The different detected species of C_L_-TMD constructs are indicated with arrows. Percentages below the blot indicate the C-terminally glycosylated fraction, giving an estimate of the fraction that fails to integrate into the ER membrane. The asterisk (*) denotes a fraction of cleaved species. Cleavage was also previously observed for similar reporter constructs containing the C_L_ domain, which occurred post-lysis (Behnke *et al*, 2016). **(B)** Predicted ΔG_app_ values for membrane integration for the isolated TMDs of investigated Cx32 mutants. **(C)** EndoH digest of the C_L_-TMD3 domain reporter and its indicated mutants to assess their membrane integration. Percentages below the blot indicate the C-terminally glycosylated fraction, giving an estimate of the fraction that fails to integrate into the ER membrane. **(D)** Illustration of the amino acid sequence and construct details of minCNX and **(E)** of minCNX^ΔTMD^.

**Appendix Fig S3.**
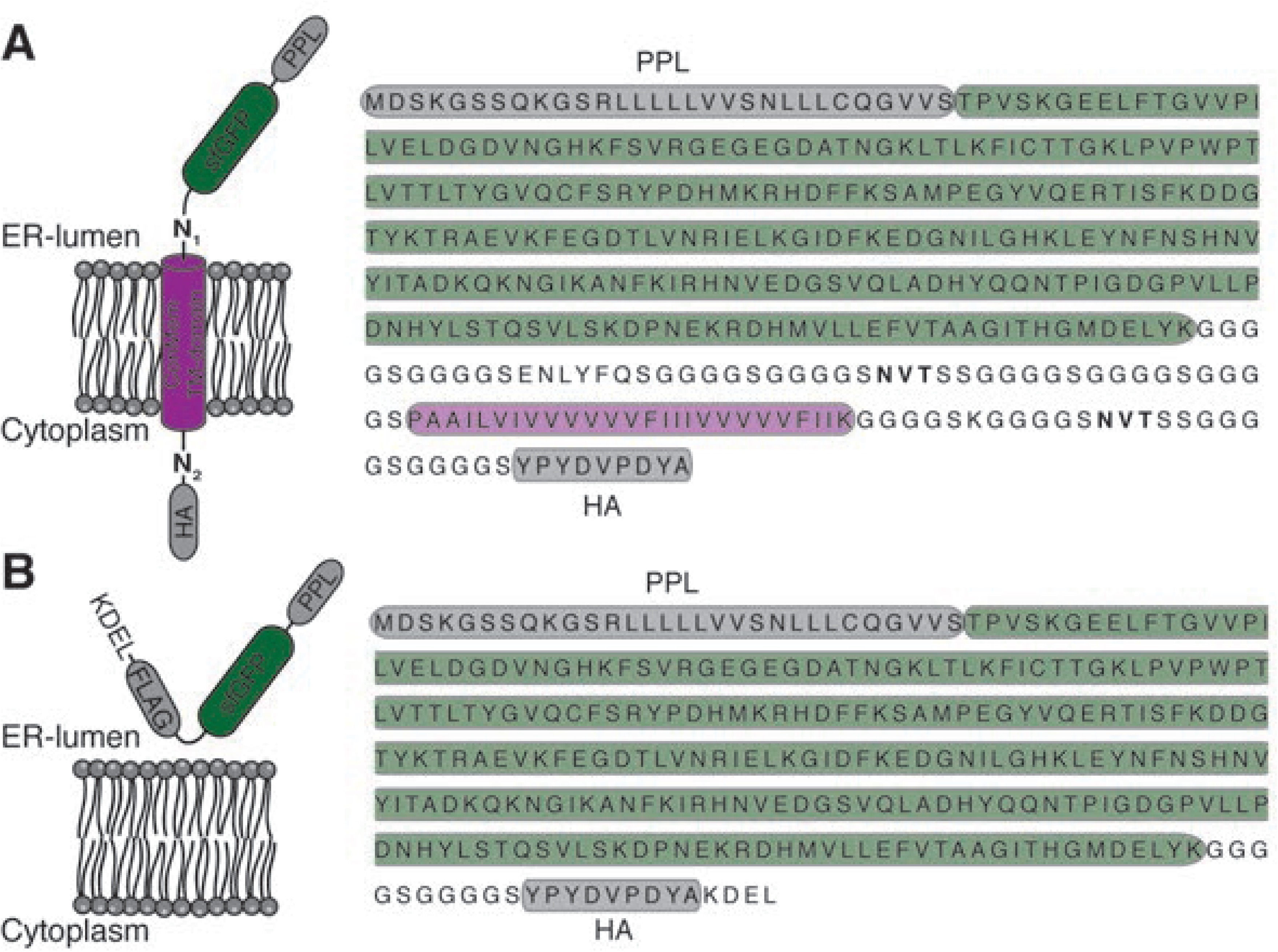
Schematic of the ConMem and ConMem^ΔTMD^ construct. (**A**) Illustration of the amino acid sequence and construct details of ConMem and **b,** of ConMem^ΔTMD^. To further promote a type I orientation of ConMem, an additional Lys residue (in addition to the C-terminal one in the consensus sequence) was placed between the TMD and the NVT reporter site according to the positive inside rule (von Heijne & Gavel, 1988).

**Appendix Fig S4.**
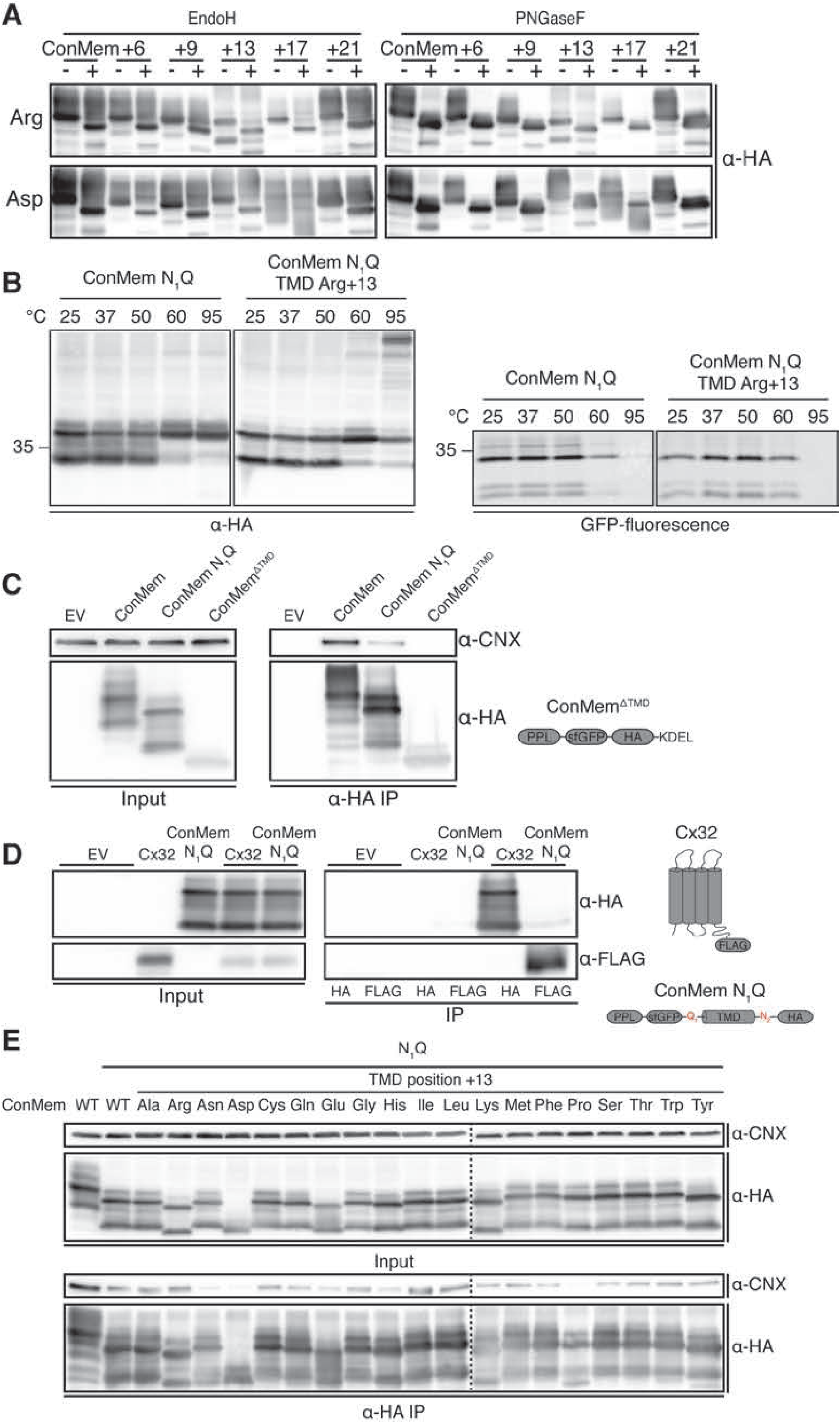
Design and validation of ConMem. **(A)** ConMem and ConMem mutant variants containing amino acid replacements against arginine and asparagine at the indicated positions of the TMD region were transfected into HEK293T cells. Cell lysates were treated with or without EndoH or PNGaseF as indicated and analyzed by immunoblotting. Comparison of the glycosylation pattern with respect to ConMem indicates no misintegration for the different ConMem TMD mutants. **(B)** ConMem N_1_Q and ConMem N_1_Q Arg+13 were transfected into HEK293T cells. Cell lysates were split and exposed to different denaturing temperatures in Laemmli buffer as indicated. Analysis by immunoblotting and fluorescent imaging of sfGFP revealed some aggregation at higher temperatures. The two lower species correspond to ConMem N_1_Q with an unfolded (upper) or a folded (lower, more compact) sfGFP moiety, as can be seen in the fluorescence-imaged gel on the right to detect sfGFP fluorescence. **(C)** CNX does not bind to ConMem^ΔTMD^. HEK293T cells were transiently transfected with ConMem, ConMem N_1_Q or ConMem^ΔTMD^ as indicated. Cell lysates and HA-immunoprecipitates were analyzed for HA-tagged ConMem variants and co-immunoprecipitating endogenous CNX. **(D)** Cx32 does not interact with ConMem. HEK293T cells were transiently transfected with either HA-tagged ConMem N_1_Q constructs, FLAG-tagged Cx32 constructs or both together. HA-Immonoprecipitates were analyzed for ConMem N_1_Q and co-immunoprecipitating Cx32 and vice-versa. **(E)** Representative blots of co-immunoprecipitation experiments between ConMem and CNX from HEK293T cells transfected with the indicated ConMem constructs, each of which contained one of the 20 amino acids on position 13 within the TMD region.

**Appendix Fig S5.**
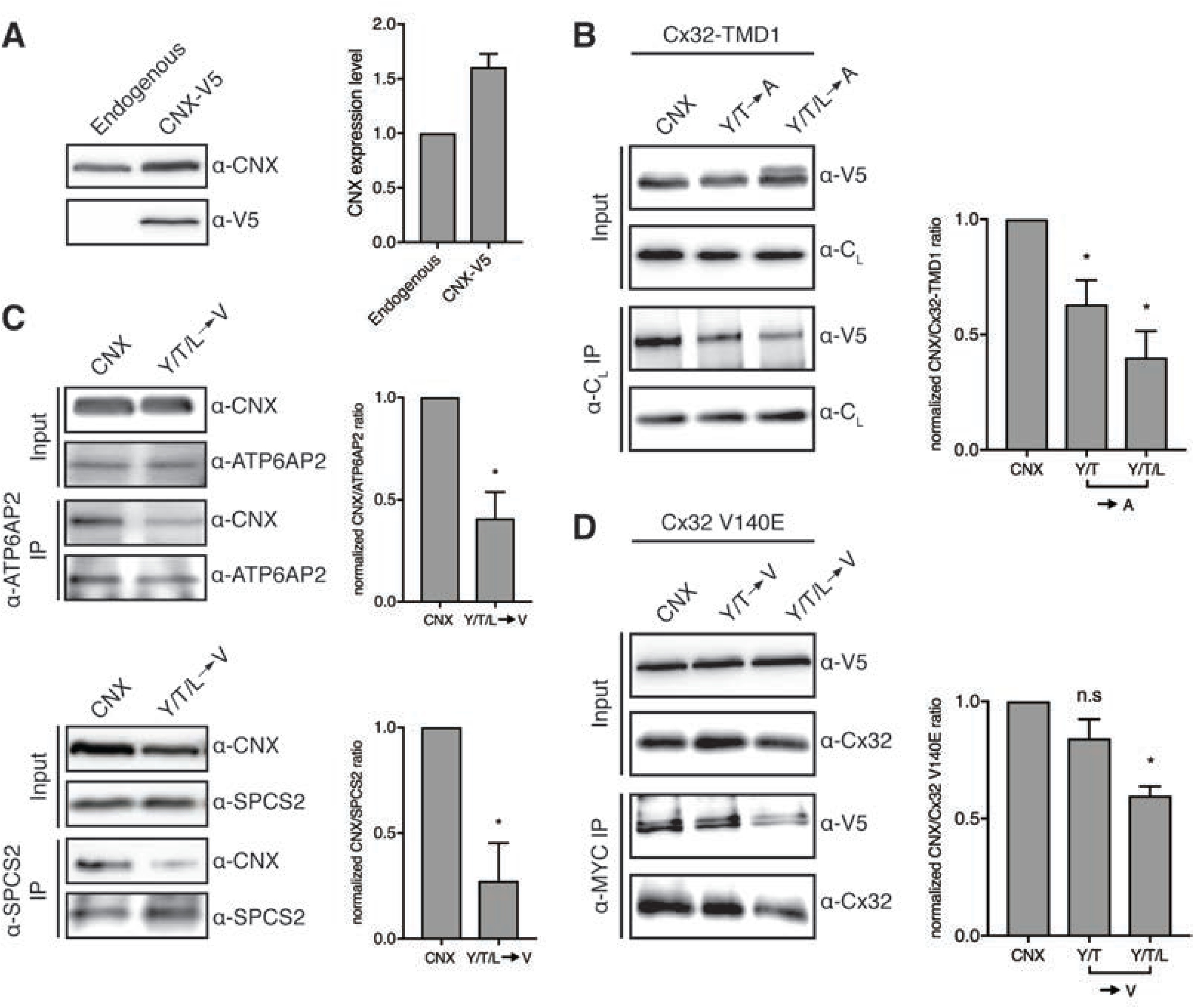
Client interaction analyses for CNX mutants. **(A)** Overall CNX level in CNX-V5 expressing cells show approximately 1.5-fold higher CNX levels compared to endogenous CNX expression levels. Cell lysates of untransfected and HEK239T cells transiently transfected with CNX-V5 constructs were immunoblotted against CNX and against the V5-tag. One representative blot is shown with quantifications on the right (mean ± SEM, N ≥ 3, *P< 0.05, two-tailed Student’s t tests). **(B)** Interaction of Cx32-TMD1 is equally reduced for CNX mutants where amino acids important for interaction as shown in Fig. 5A were exchanged to alanine instead of valine residues. HEK293T cells were transiently transfected with the indicated constructs and cell lysates and Cx32 TMD1 immunoprecipitates were analyzed for C_L_-tagged Cx32 TMD1 and co-immunoprecipitating CNX mutants. One representative blot is shown with quantifications on the right (mean ± SEM, N ≥ 5, *P< 0.05, two-tailed Student’s t tests). **(C)** Interaction of the single-pass TM protein ATP6AP2 and the dual-pass TM protein SPCS2 with V5-tagged CNX. CNX knockout HEK293T cells were transiently transfected with the indicated V5-tagged CNX constructs and cell lysates and ATP6AP2 or SPCS2-immunoprecipitates were analyzed for endogenous ATP6AP2 or SPCS2 and co-immunoprecipitating CNX. One representative blot is shown. In each case, the graph shows results from at least four independent experiments (mean ± SEM, N ≥ 4, *P value < 0.05, two-tailed Student’s t tests). **(D)** Interaction of Cx32 V140E full-length with CNX behaves similarly compared to Cx32 WT. HEK293T cells were transiently transfected with the indicated constructs. Cell lysates and MYC-tagged Cx32 V140E immunoprecipitates were analyzed for Cx32 V140E and co-immunoprecipitating CNX mutants. One representative blot is shown. The graph shows results from six independent experiments (mean ± SEM, N ≥ 6, n.s.: not significant, *P value < 0.05, two-tailed Student’s t tests).

**Appendix Fig S6.**
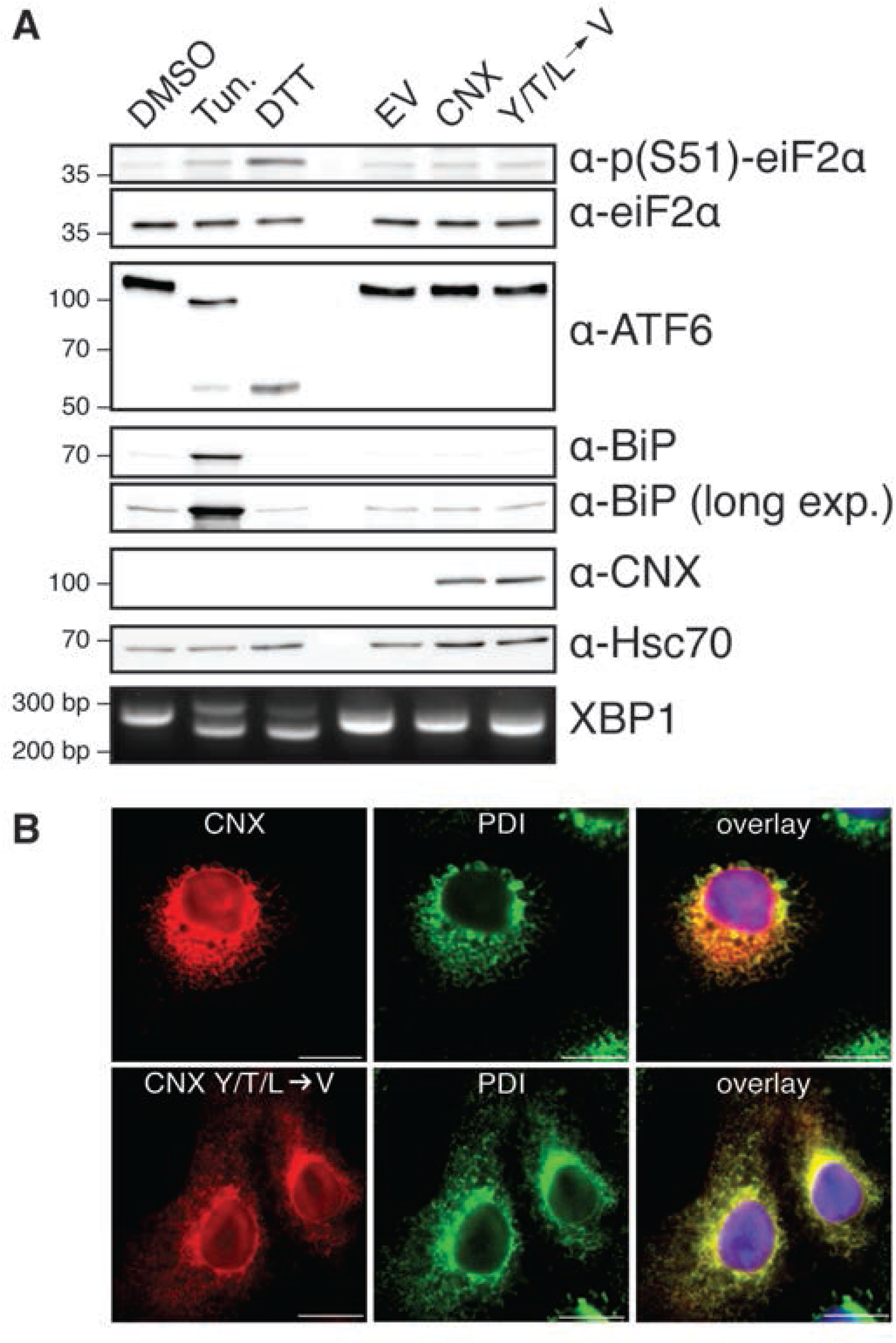
Validation of the HEK293T CNX knockout cell line for functional assays. **(A)** UPR activation was assessed in CNX knockout cells transfected with mock plasmid, and incubated with DMSO (vehicle control), tunicamycin (Tun.), or DTT. CNX, or its binding-deficient mutant were transfected into the CNX knockout cells (right side) and activation of the the same branches of the UPR was assessed. **(B)** COS-7 cells were transfected with the indicated CNX ALFA-tagged constructs and fluorescence microscopy was performed using anti PDI (green) as an ER marker. CNX WT constructs and CNX mutant variants deficient in substrate binding were detected with anti ALFA-FluoTag-X2 nanobodies (red) and show ER localization. Nuclei were stained with DAPI (blue). Images are representative of cells from at least three different biological replicates. Scale bars correspond to 20 µm.

**Appendix Fig S7.**
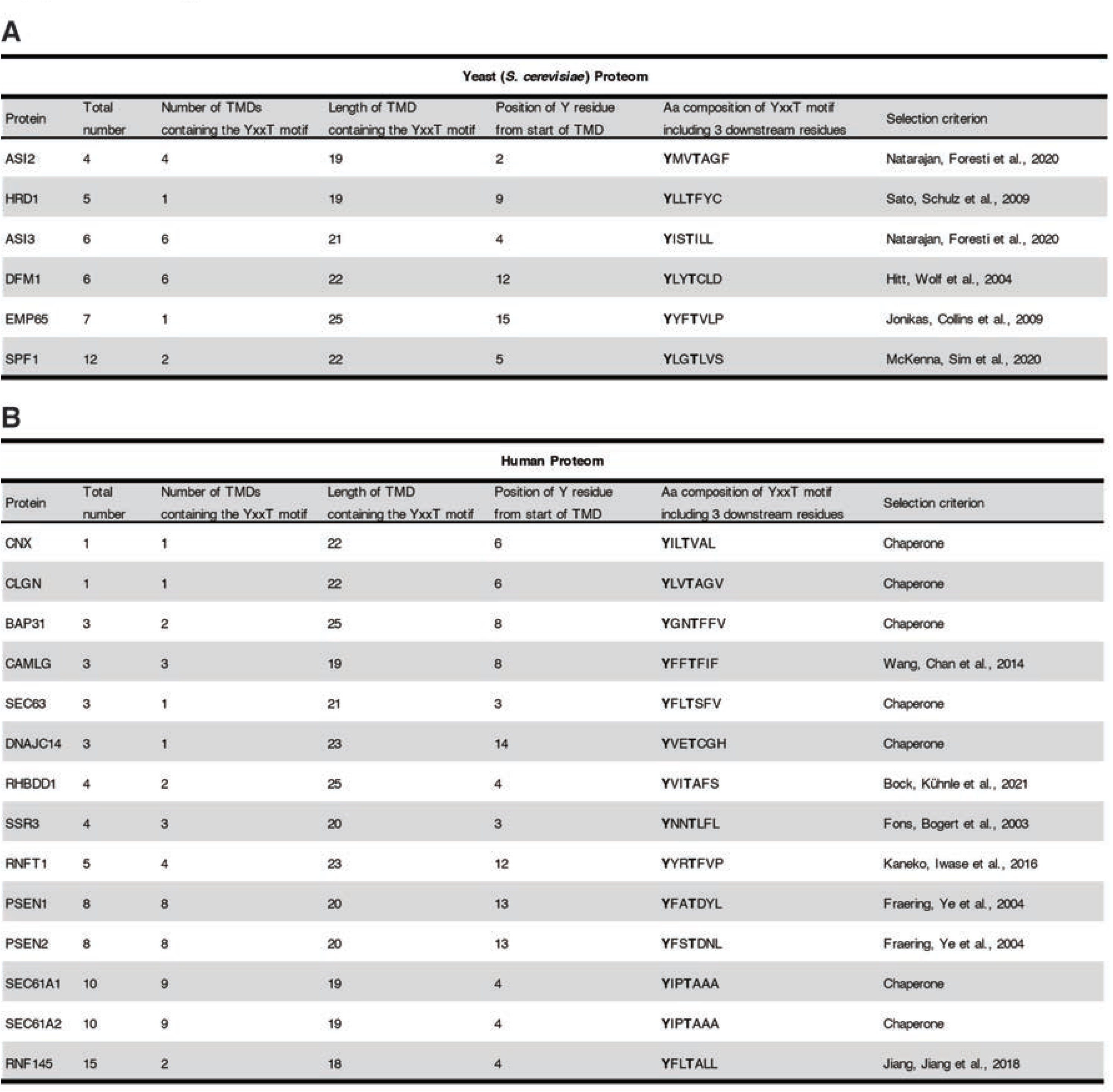
Further analyses of the YxxT motif in Calnexin and other proteins. Search for YxxT motifs in all ER/inner nuclear membrane proteins in **(A),** yeast and **(B),** humans. TM regions were predicted using Phobius and analysis for YxxT motifs was performed from 3 residues downstream of the N-terminus of the TM region until 3 residues upstream of the C-terminus, resulting in an overall amount of 43 hits for the yeast and 103 hits for the human proteom. From those hits, listed are the gene names of the respective proteins, along with additional information regarding the TMD and motif such as sequence, that are either annotated in Uniprot as chaperones or have other published functions related to intramembrane quality control. In the latter case, one reference for their respective function is given in the selection criterion column.

---

**General:**

CNX HEK293T CNX KO cells transfected with vector coding for Calnexin

eV HEK293T CNX KO cells transfected with empty vector as control

**CNX enrichment filtered:**

Tables were exported from Perseus after full processing. Data shown are filtered for p-value [-log10] ≥ 1.3 and Ratio [log2] CNX /eV ≥ 1.

1-3 number of replicates/ clones

**Proteome analysis:**

Tables were exported from Perseus after full processing. Data shown are filtered for p-value [-log10] ≥ 1.3 and Ratio [log2] eV / CNX ≤ 0.27 1-4 number of replicates/ clones

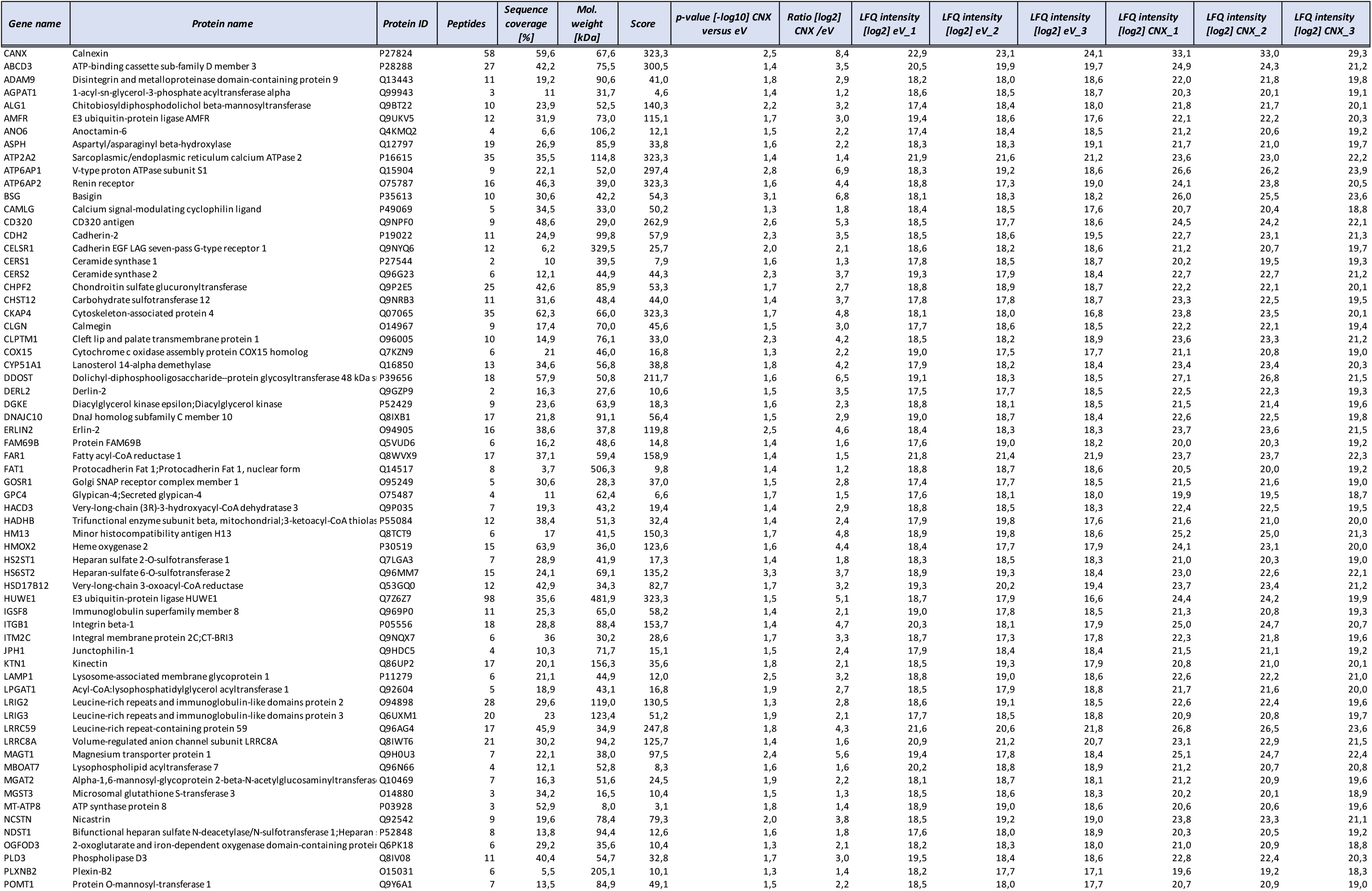
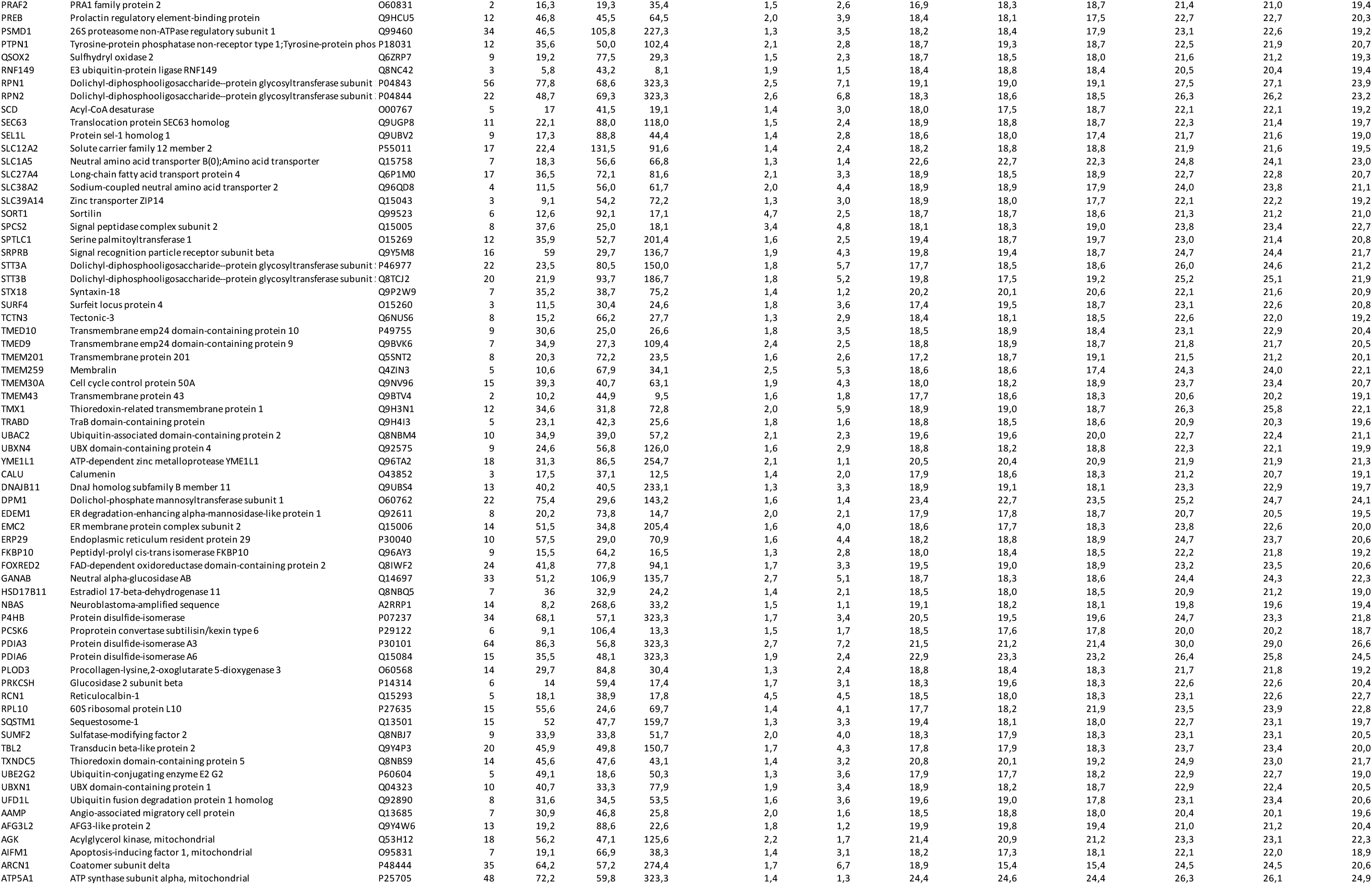
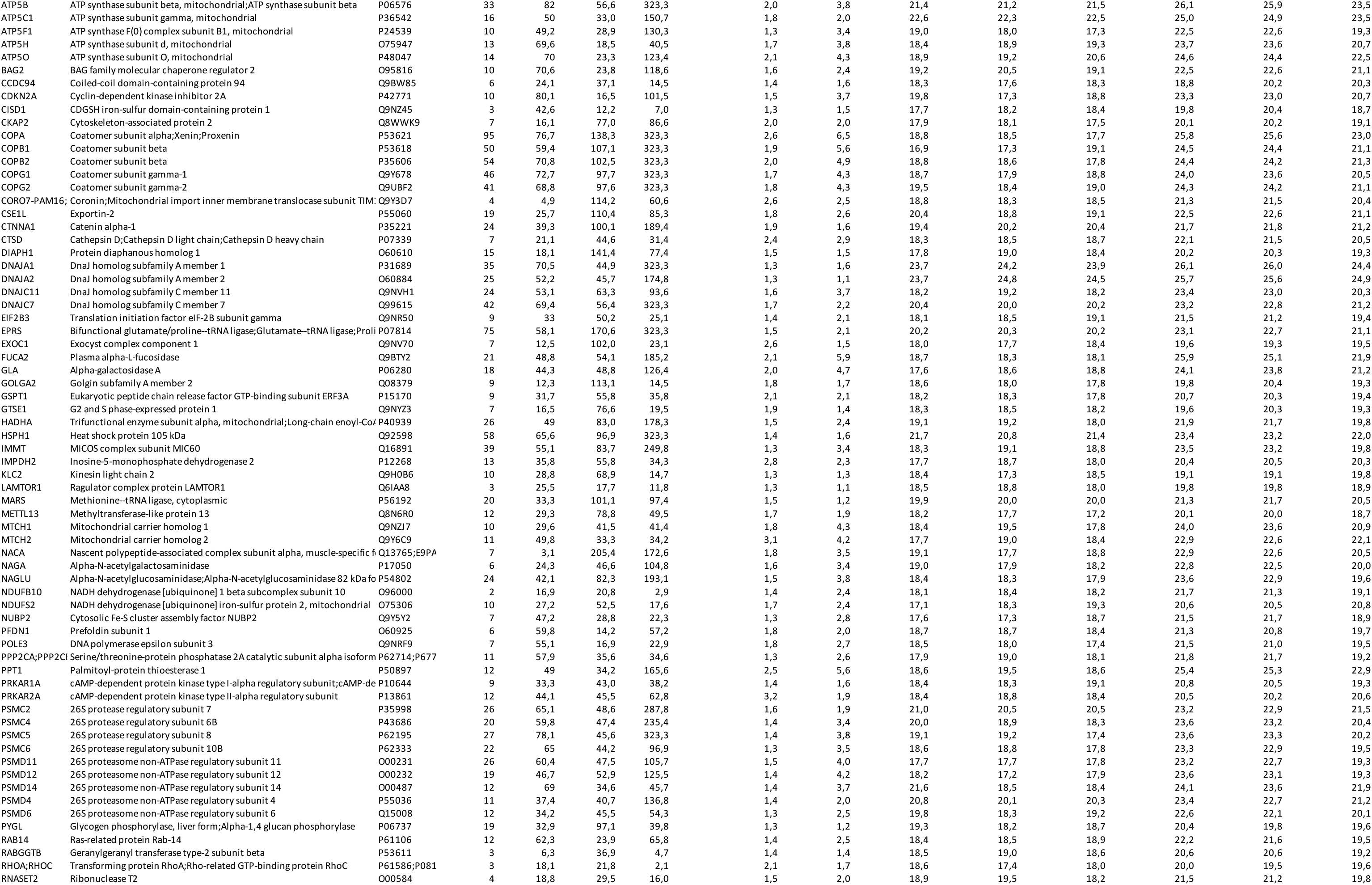
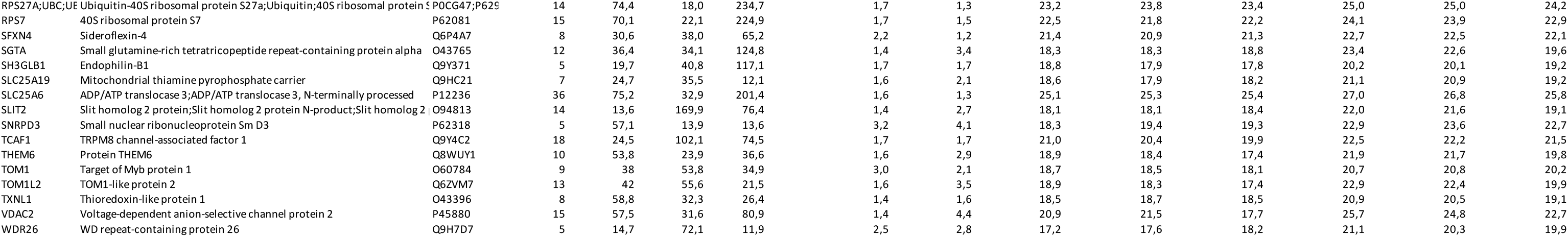

**General:**

CNX HEK293T CNX KO cells transfected with vector coding for Calnexin

eV HEK293T CNX KO cells transfected with empty vector as control

**CNX enrichment filtered:**

Tables were exported from Perseus after full processing. Data shown are filtered for p-value [-log10] ≥ 1.3 and Ratio [log2] CNX /eV ≥ 1.

1-3 number of replicates/ clones

**Proteome analysis:**

Tables were exported from Perseus after full processing. Data shown are filtered for p-value [-log10] ≥ 1.3 and Ratio [log2] eV / CNX ≤ 0.27 1-4 number of replicates/ clones

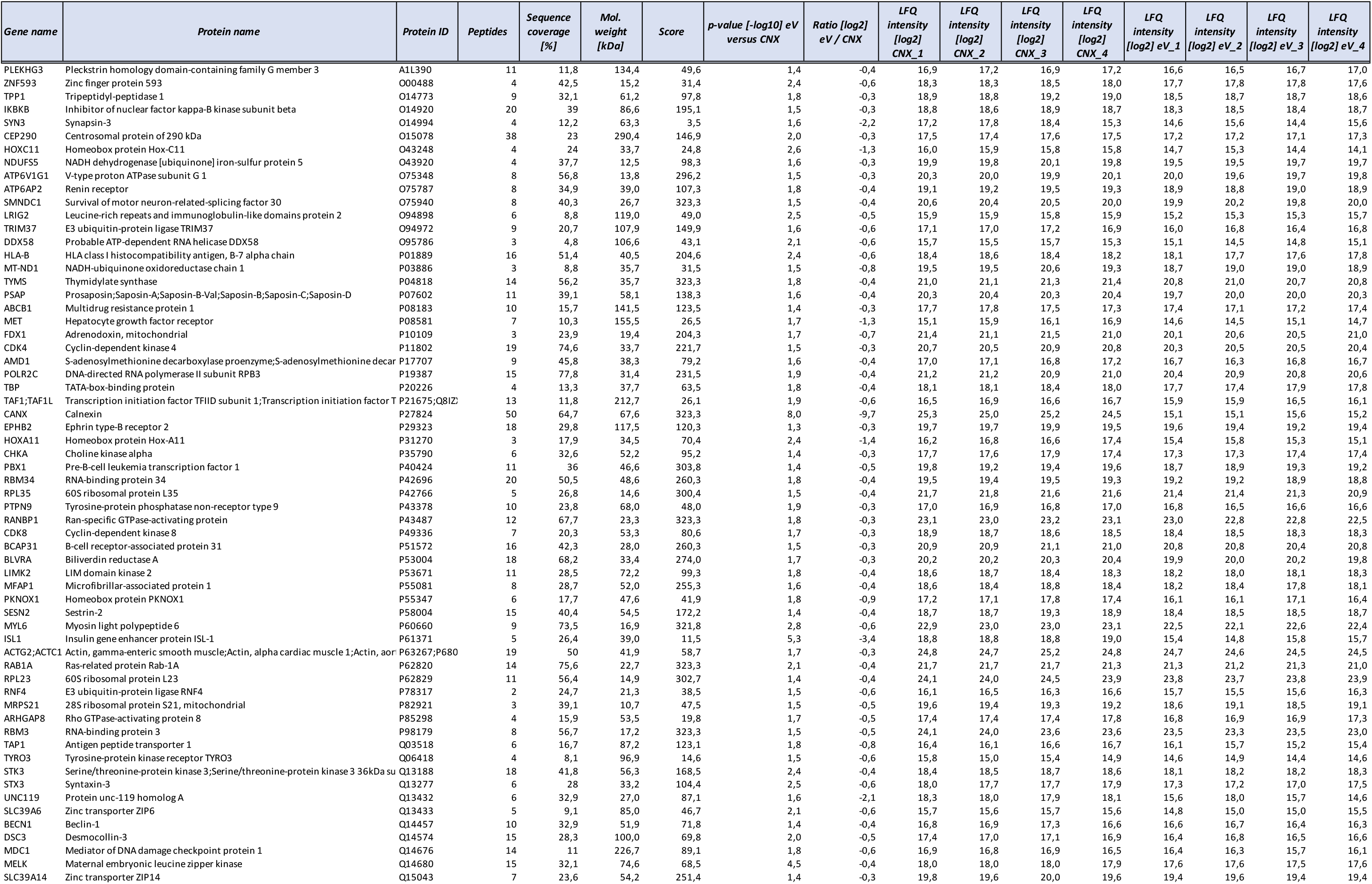
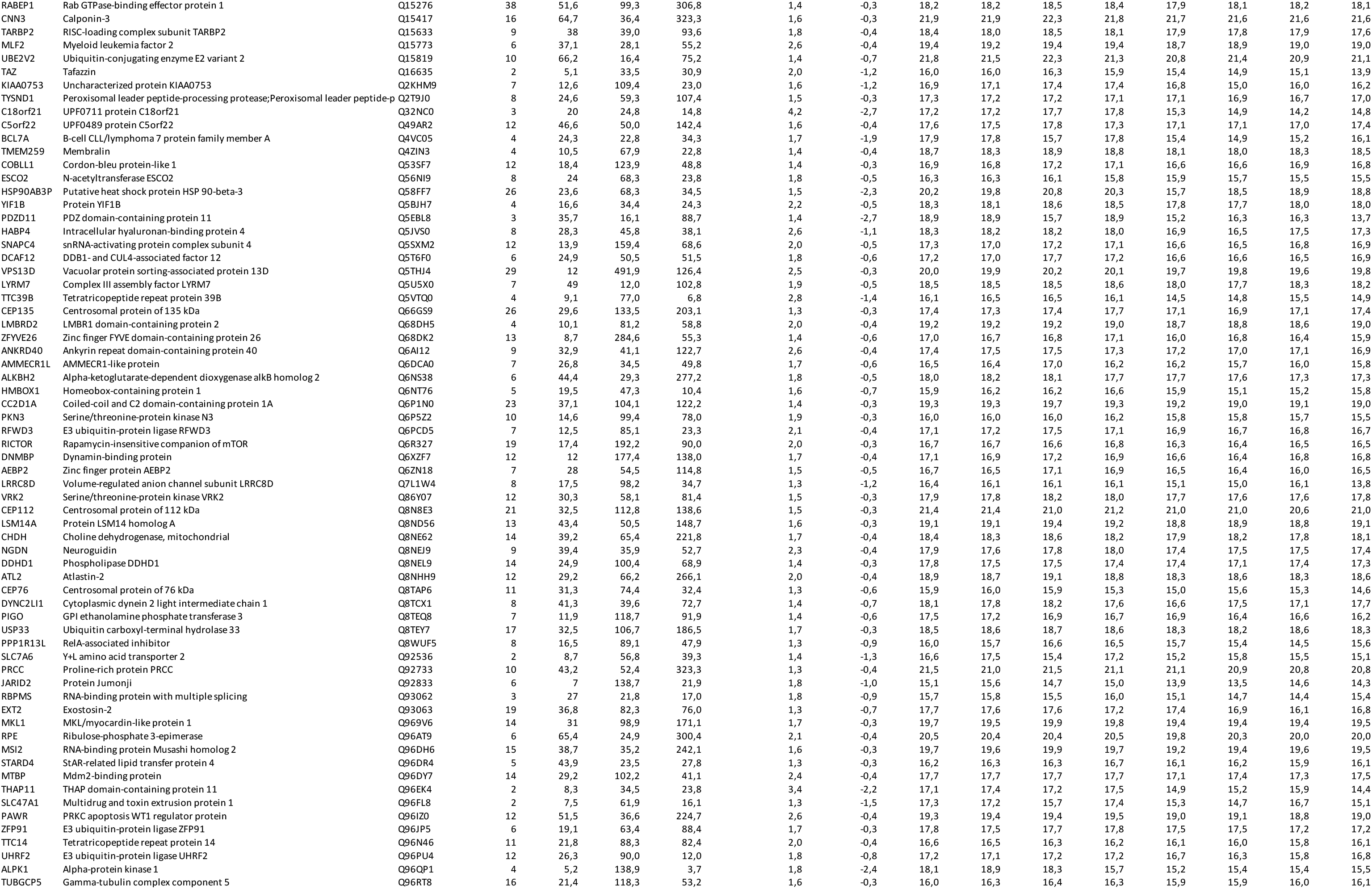
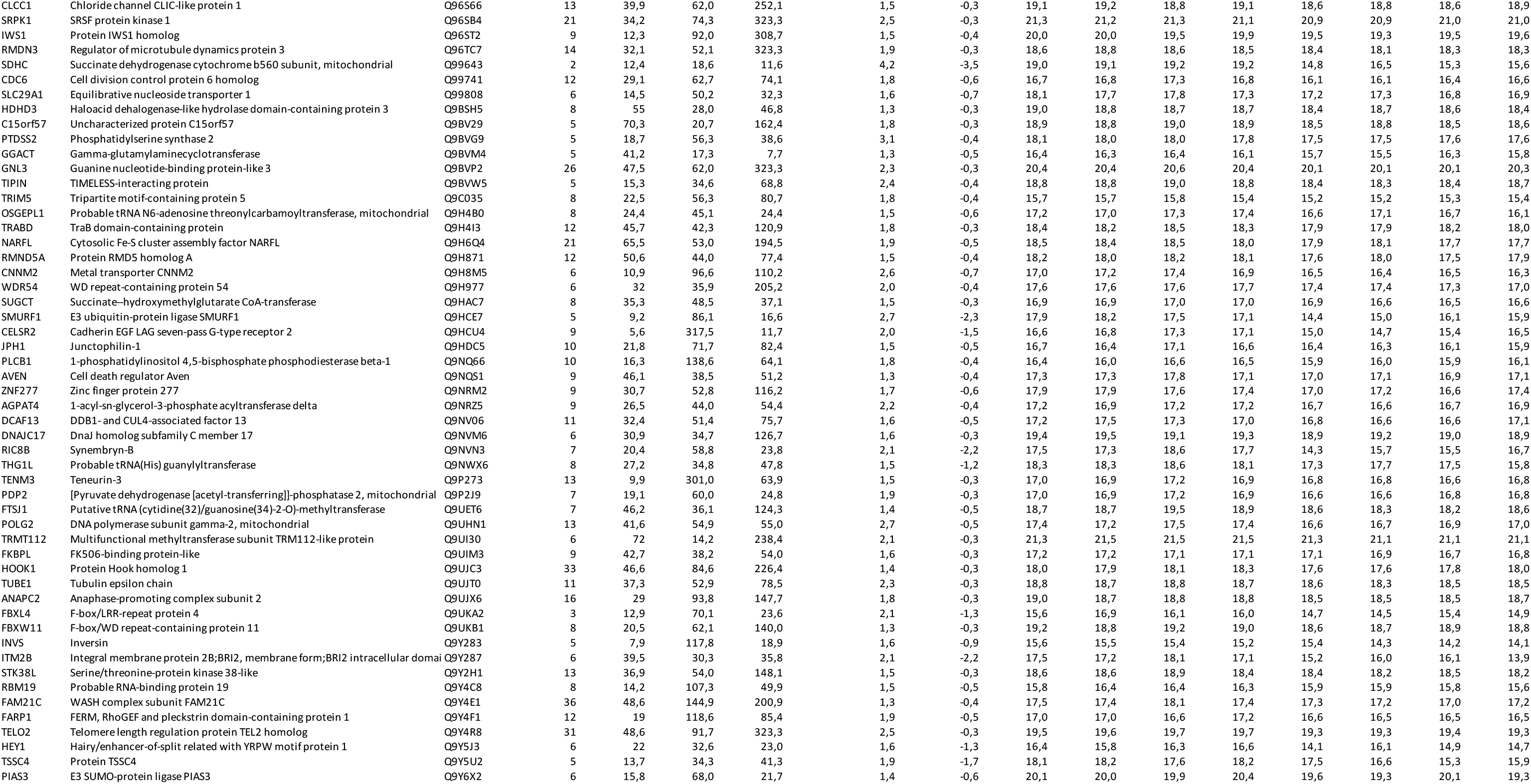

